# The UBR box E3 ligases Poe and Hyd are required for efficient Pericentrin degradation

**DOI:** 10.1101/2022.01.21.477277

**Authors:** Ramya Varadarajan, Brian J. Galletta, Carey J. Fagerstrom, Karen Plevock Haase, Nasser M. Rusan

**Affiliations:** Cell and Developmental Biology Center, National Heart Lung and Blood Institute, National Institutes of Health, Bethesda, MD

## Abstract

Centrosomes are the major microtubule organizing centers (MTOC) of many cells and are composed of centrioles and the pericentriolar material (PCM). Centrosomes are an essential part of diverse cellular processes and require precise regulation of their protein levels. One protein whose levels must be regulated is Pericentrin – PCNT in humans and PLP in *Drosophila*. Increased PCNT expression and its protein accumulation is linked to clinical conditions including cancer, mental disorders, and ciliopathies. However, the mechanisms by which PCNT levels are regulated remains underexplored. Our previous study (Galletta, 2020) demonstrated that PLP levels are sharply downregulated during early spermatogenesis and this regulation is essential to spatially position PLP on the proximal end of centrioles. We hypothesized that the sharp drop in PLP protein was a result of a rapid protein degradation during the male pre-meiotic G2 phase. In this study we show that the N-terminal region of PLP is required for regulating its levels, which when elevated, extends PLP’s position distally on centrioles. Consequently, PCM was also mispositioned leading to defects in spermatids. We then performed a targeted screen where we identified the E2 ligases, Rad6 and UbcD1, and the UBR box family E3 ligases, Poe (UBR4) and Hyd (UBR5) as factors that promote PLP degradation in spermatocytes. Collectively, our study suggests that Poe and Hyd directly bind the N-terminus of PLP and target it for degradation, ensuring proper centriole/basal body assembly and male fertility.

## Introduction

Centrosomes, composed of centrioles and the surrounding pericentriolar material (PCM), are the major microtubule organizing centers in many cells. They serve critical functions including organizing and orienting spindles during cell division, ensuring proper docking of the sperm tail to the nucleus during spermiogenesis and many more (Avidor-Reiss, 2019; Galletta, 2020; Hoffmann, 2021; Wu, 2020; Yuan, 2015). The proper regulation of the assembly, activity, and disassembly of centrosomes is critical for cellular and tissue function and, as such, for proper organismal development. Defects in these processes can arise through the presence of too few or too many centrosomes, through inactive or over-active centrosomes, and even by changes in sub-organellar structure. Therefore, centrosomes must be tightly regulated to ensure they are intact and functional at the correct time and place (Nigg, 2018; Ryniawec, 2021; Schatten, 2018).

Significant study of the regulation of centrosomes has focused on regulation of protein-protein interaction, much of which relies on kinases triggering a phosphorylation event that creates or disrupts a protein interaction surface (Boese, 2018; Conduit, 2014; Dzhindzhev, 2014; Kratz, 2015; Ohta, 2014; Woodruff, 2015; Wueseke, 2016). Another critical aspect of centrosome protein regulation is ensuring their correct levels at the centrosome or globally throughout the cell (Arquint, 2014; Arquint, 2012; D’Angiolella, 2010; Keller, 2014; Strnad, 2007) in coordination with the cell cycle. Global protein level regulation is particularly interesting and complex as it could involve multiple mechanisms. For any protein with oscillatory levels, like one synchronized with the cell cycle, one could postulate regulation at the level of transcription, mRNA stability, translation, or post-translation. A major focus in the area of protein level control has been on mechanisms of protein degradation, which is critical for proper centrosome form and function (Badarudeen, 2021; Zhang, 2016). Inhibition of the proteasome leads to overduplication of the centriole (Duensing, 2007), centriole elongation (Korzeniewski, 2010), and an expansion in the PCM (Wigley, 1999). More targeted studies in recent years have uncovered many protein degradation mechanisms that control centrosome protein levels both at the centrosome and throughout the cell, including the major centrosome regulatory kinases Plk4/ZYG-1 (Cajanek, 2015; Cunha-Ferreira, 2013; Holland, 2010; Medley, 2021; Peel, 2012; Rogers, 2009) and Polo/Plk1 (Braun, 2021; Fang, 1998; Lindon, 2004), and centriole proteins such as CP110 (D’Angiolella, 2010), STIL/Ana2/Sas-5 (Arquint, 2014; Arquint, 2012; Medley, 2017; Tang, 2011), CPAP/Sas4 (Tang, 2009) and Sas-6 (Badarudeen, 2021; Badarudeen, 2020; Puklowski, 2011; Strnad, 2007). This suggests a tight, multilevel regulation of not only protein activity, but also protein availability throughout the cell.

Most of the published protein degradation studies relate to centriole duplication control; considerably less is known about the regulation of PCM proteins by protein degradation. One example is the *Drosophila* PCM protein Spd2, global levels of which are under the control of APC/C^Fzr^ (Meghini, 2016). The mammalian ortholog of Spd2, Cep192, is also under the control of protein degradation, via SCF^FBXL13^ (Fung, 2018). Interestingly, modulating the levels of SCF^FBXL13^ not only tunes the level of Cep192 at the centrosome, but also affects the recruitment of gamma-tubulin and the MTOC activity (Fung, 2018). It is therefore likely that regulation of other PCM proteins globally is used to regulate MTOC function in the context of the cell cycle or in other cellular circumstances.

The *Drosophila* male germline provides an excellent model for studying the assembly of centrosomes and the regulation of its component proteins. Germ cells undergo a series of amplifications to produce spermatocytes (SCs) that then enter a prolonged G2 phase. This phase has served as an important model for studying centriole elongation as they elongate to ∼1.8 microns, 10 times their starting length. It is during this period that distinct organizations of proteins along the proximal and distal axis of the centriole are established. The proximal end of the centriole organizes PCM and interfering with the organization of proximal PCM disrupts the construction of functional sperm at later stages of spermatogenesis (Galletta, 2020). This proximal position of PCM is dictated by the position of the Pericentrin-like Protein (PLP), a member of the “bridge” protein class that localize to the centriole wall and extend into the PCM as a scaffold and a regulator of PCM (Varadarajan, 2018). The position of PLP itself along the centriole is dictated by the timing of its cellular availability during centriole elongation. At the beginning of centriole elongation in SCs, PLP mRNA and protein are available, but as centriole elongation proceeds both are no longer available, resulting in a restriction of PLP to the oldest, most proximal, portions of the centriole (Figure 1A; Galletta, 2020). Experimentally forcing PLP availability later in G2 results in PLP occupying more distal positions on the centriole, PCM accumulating in improper distal positions, and ultimately defects in the docking of the centriole to the nucleus in spermatids yielding decapitated sperm (Galletta, 2020). The importance of regulating the levels of PLP does not appear to be limited to flies, as an increase in the levels of human Pericentrin mRNA and protein by just 50%, as occurs in individuals with trisomy 21, is sufficient to cause cilia defects (Galati, 2018).

**Figure 1:**
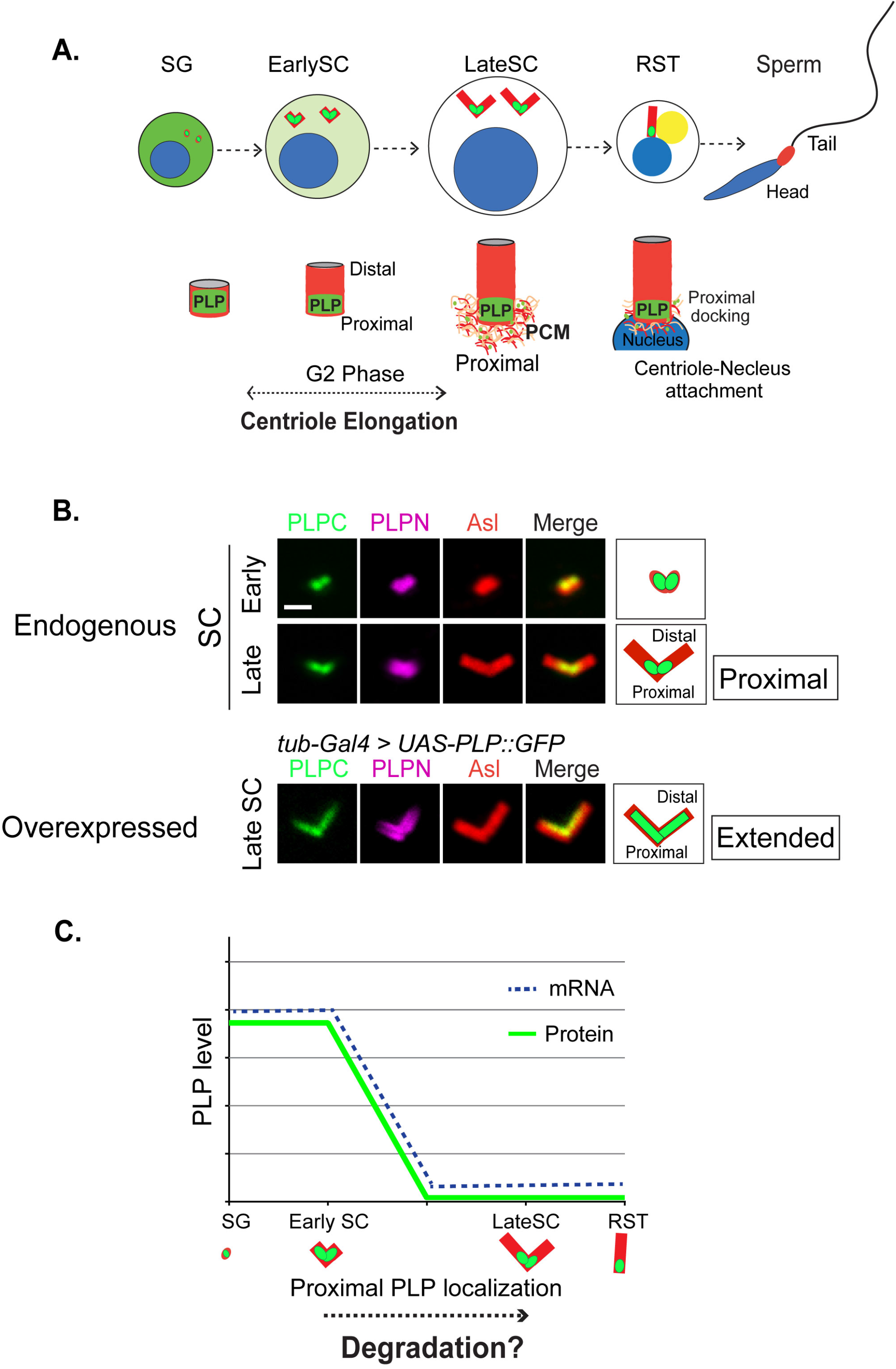
PLP levels are downregulated in developing spermatocytes. **(A)** Cartoon representation of centriole elongation during G2 of spermatocyte development showing cytoplasmic and centriole levels of PLP (Green). PLP remains proximal and dictates spatial recruitment of PCM. Centriole (Red), Nucleus (Blue), Mitochondrial Derivative (Nebenkern; Yellow), Spermatogonia (SG), Spermatocytes (SC), Round spermatids (RST). **(B)** Proximal localization of endogenous PLP in SC shown using CRISPR knock-in mNeon tag on the C-terminus (PLPC, green) and N-terminal antibody (PLPN, magenta). Centriole labeled by antibody to Asl (red). *tubulin-Gal4>UAS-PLP::GFP* is localized along the full length of the centrioles. Bar = 1 µm. **(C)** Representation of PLP protein level (Green), mRNA availability (Blue) during spermatogenesis.

While some of the PLP protein level regulation is the result of the timing of transcript availability, we hypothesized that additional regulation at the protein level is also required to precisely control the timing of PLP availability during germline development, especially to ensure that its levels dramatically drop as the pre-meiotic G2 proceeds, and centrioles elongate. In this study we demonstrate that PLP is regulated by post-translational protein degradation in the germline. We find that specific regions of PLP are critical for this regulation, and that several E2 and E3 ligases are involved. PLP is therefore under multiple levels of regulation to ensure the proper timing of its availability during centriole elongation.

## Results

### The N-terminus of PLP restricts its position to the proximal end of centrioles

Our previous work established that the proximal position of PLP on spermatocyte centrioles is dictated by the timing of available PLP protein (Figure 1A-C; Galletta, 2020). PLP is available when centrioles are initially built in spermatogonia (SG) and early spermatocytes (SC), but not available as the centrioles elongate in the premeiotic G2 phase SCs (Figure 1A-C). During G2 both the levels of *PLP* mRNA and PLP protein drop significantly, indicating that PLP is tightly regulated at multiple levels (Figure 1C; Galletta, 2020). Consistent with our previous observations, increasing PLP protein availability by driving expression under the control of a ubiquitous promoter (*ubi-PLP::GFP*), or *tubulin-Gal4* >*UAS-PLP::GFP*, results in PLP localizing along the entire centriole length (Figure 1B) and accumulating to higher levels in the tissue (Figure S1A).

The importance of properly regulating PLP protein levels led to the hypothesis that PLP is under strict post-translational regulation via degradation (Figure 1C). We began to investigate this hypothesis by performing a structure-function analysis to identify regions of PLP potentially required for regulating PLP levels. We generated a series of transgenic animals expressing N-terminal PLP deletions under the control of the UAS promoter; all transgenes maintained the C-terminal centriole targeting motif (PACT) intact (Figure 2A). To control the timing of transgene expression, we used the *bam-Gal4* driver, which expresses in early germline development prior to centriole elongation. We found that full length *PLP* (*PLP^FL^*) driven by *bam*-*Gal4* localized only to the centriole proximal end (Figure 2B; Galletta, 2020), similar to the endogenous protein. This indicated that the timing of *PLP* mRNA expressed by the *bam* promoter was sufficiently similar to the endogenous *PLP* promoter to result in grossly normal centriole localization. Unless otherwise noted, all subsequent experiments were performed using UAS transgenes driven by *bam-Gal4*.

**Figure 2:**
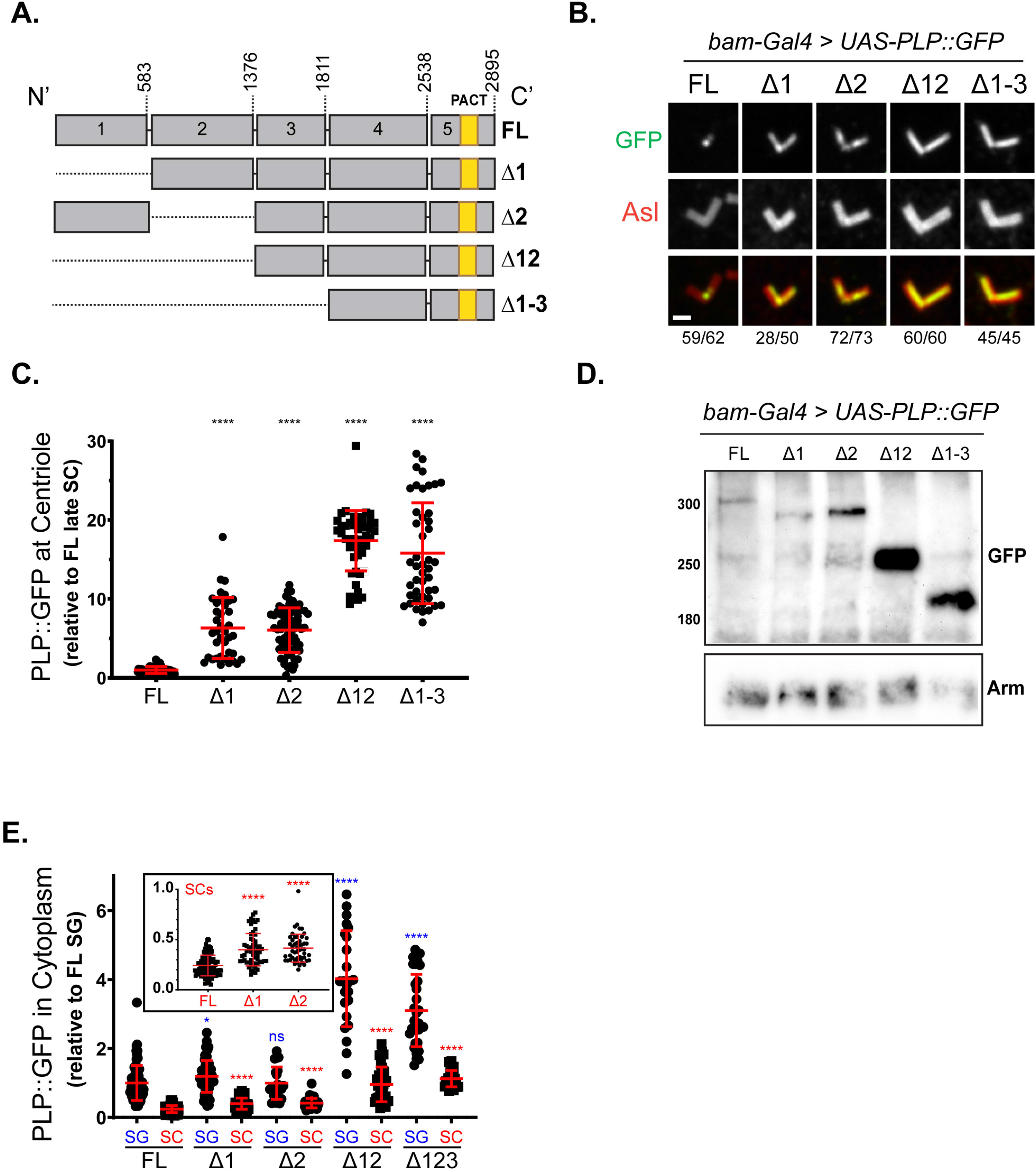
The N-terminus of PLP regulates its levels and centriole position. **(A)** Schematics of N-terminal PLP deletion constructs. Grey regions represent protein fragments 1 through 5, the PACT domain is indicated as a yellow bar. Full length (FL) represents the PLP-F isoform. Δ1 = delta aa1-583, Δ2 = delta aa584-1376, Δ12 = delta aa1-1376, and Δ1-3 = delta aa1-1811. **(B)** Late SC centrioles from flies expressing *bam-Gal4>UAS-PLP::GFP* (green) constructs. Centriole labeled by antibody to Asl (red). Fraction of centriole pairs showing these localization patterns are indicated below each column. At least 9 testes of each genotype were examined. All deletion constructs show extension of PLP along the entire centriole length. Additional localizations are shown in Supplemental Figure 1B. Bar = 1 µm. **(C)** Quantification of the fluorescent intensity of PLP::GFP on centrioles in late SCs. The measurements are shown relative to the PLP-FL. FL (n = 62), Δ1 (n = 38), Δ2 (n = 72), Δ12 (n = 44), Δ1 -3 (n = 45). At least 9 testes of each genotype were examined. **(D)** Western blot of testes from indicated genotypes. Anti-GFP antibodies were used to detect PLP and anti-Armadillo was used as a loading control. **(E)** Cytoplasmic PLP::GFP level in spermatogonia (SG) and spermatocytes (SC) in testes expressing *bam-Gal4>UAS-PLP::GFP* transgenes. Measurements are shown relative to the cytoplasmic level of PLP-FL in SG. Cells analyzed - FL (SG n = 73; SC n =107), Δ1 (SG n = 107; SC n = 57), Δ2 (SG n = 21; SC n =51), Δ12 (SG n = 24; SC n = 38), Δ1-3 (SG n = 33; SC n = 26). Measurements of each cell type were from at least 3 testes of indicated genotype. Statistics performed to compare SG (blue) and SC (red) separately. For quantifications, bars represent mean ± standard deviation. Statistical comparisons are to FL and by unpaired t-test with Welch’s correction when appropriate. **** = p≤0.0001, *** = p≤0.001, ns = not significant.

Deletion of either of two regions in the N-terminus of PLP (ΔF1 = delta aa1-583, ΔF2 = delta aa584-1376) resulted in PLP extending beyond the proximal centriole (Figure 2B; S1B), plus an overall elevated protein levels on centrioles in spermatocytes (Figure 2C). Co-deletion of F1 and F2 (ΔF12 = delta aa1-1376) similarly resulted in PLP localization along the entire centriole, but with an even greater increase in total protein amount (Figure 2B,C). Further N-terminal deletion (ΔF1-3 = delta aa1-1811) did not alter localization and levels beyond PLP^ΔF12^ (Figure 2B,C).

Based on the position and levels of PLP on centrioles, we predicted that cytoplasmic levels of the PLP truncations were elevated. Western blots confirmed the increase in levels of PLP^ΔF2^, and even higher levels for PLP^ΔF12^, but no further increase in PLP^ΔF1-3^ (Figure 2D), paralleling the elevated levels on centrioles. We performed a more precise measurement of cytoplasmic PLP::GFP protein using quantitative fluorescence microscopy of live SG and SC. All truncations, except for PLP^ΔF2^, were significantly elevated in SGs compared to PLP^FL^, while all were elevated in SCs (Figure 2E). Collectively these results indicate that the N-terminal region of PLP contains sequences required for regulating its own protein level. Deletion of these regulatory regions increased PLP in SG and SC cytoplasm, which in turn increased centriolar PLP levels and expanded PLP position along the distal end of centrioles.

### N-terminal PLP truncations result in reduced sperm function

Our previous work showed that overexpression of PLP resulted in the distal extension of centriolar PLP and pericentriolar material (PCM; gamma tubulin (γ−Tub)) during meiosis and in round spermatids (RSTs), which resulted in failed docking of the centriole to the nucleus and a reduction in normal sperm production (Galletta, 2020). Based on these findings, we examined the downstream consequences of the elevated PLP levels that resulted from N-terminal deletions and failed post-translational regulation. Similar to endogenous PLP, PLP^FL^ localized to the proximal end of the centriole, while the N-terminal deletions extended distally in RSTs (Figure 3A, S2A) and resulted in distal extension of γ−Tub in RSTs (Figure 3B, S2B). This redistribution of PCM via PLP resulted in erroneous, lateral docking of the centriole to the nucleus (Figure 3C, D), consistent with the model that regulation of PLP levels during the premeiotic G2 is critical for early spermatid development.

**Figure 3.**
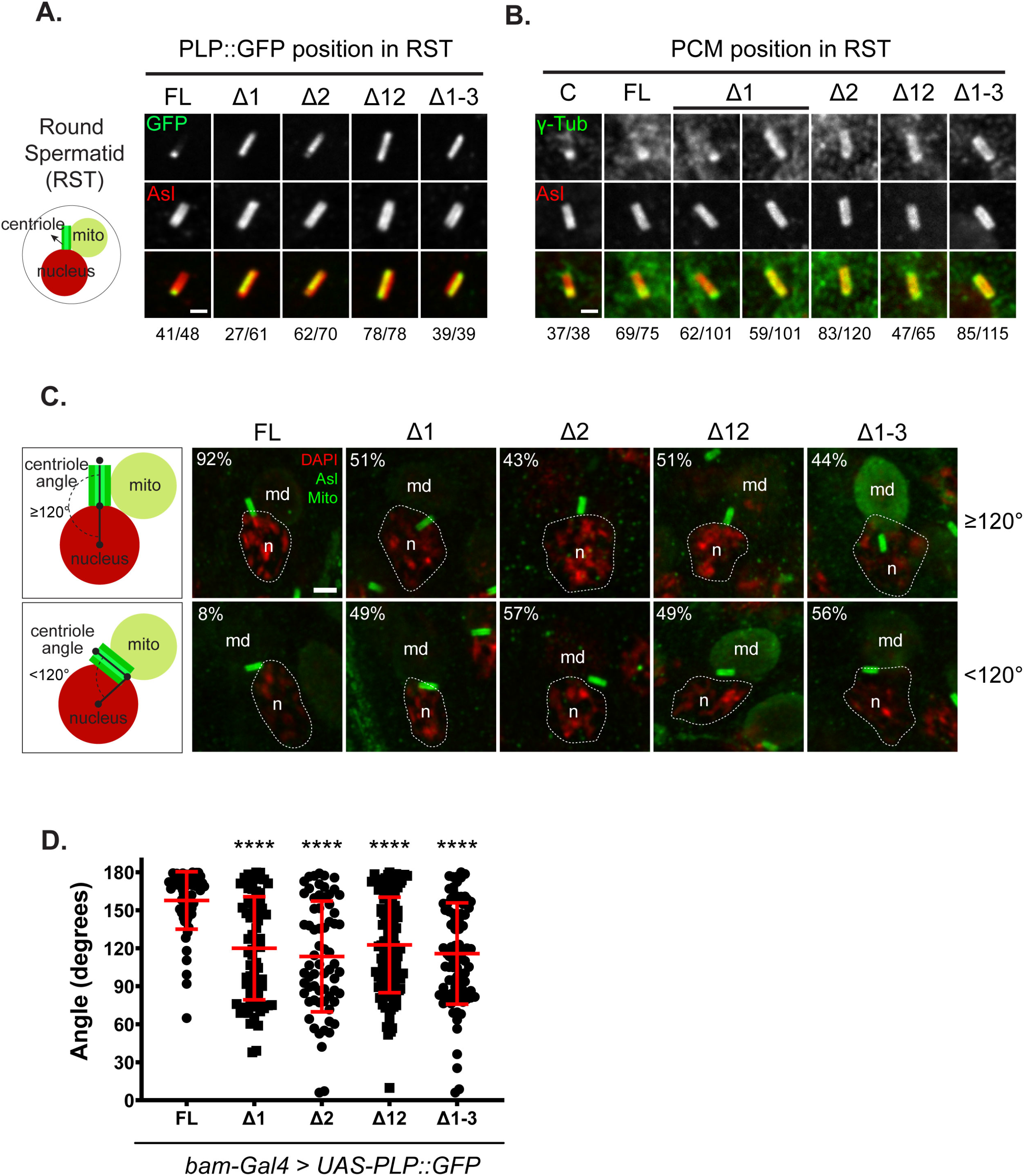
Consequences of PLP N-terminal deletions. **(A)** Centrioles from round spermatids (RSTs) from flies expressing the indicated *bam-Gal4>UAS-PLP::GFP* constructs (green). Centriole labeled by antibody to Asl (red). Fraction of centrioles showing these localization patterns are indicated below each column. At least 6 testes of each genotype were examined. All deletion constructs show extension of PLP along the entire centriole length. Additional localizations are shown in Supplemental Figure 2A. Bar = 1 µm **(B)** RST centrioles showing localization of PCM using gamma-tubulin (g-Tub, Green) on centrioles (Asl, red). Fraction of centrioles showing these PCM localization patterns are indicated below each column. At least 5 testes of each genotype were examined. g-Tub is precociously found along the entire centriole length in PLP truncations; except for PLP-Δ1 that shows PCM along the entire length only in ∼50% of centrioles. Examples of other localizations observed in Supplemental Figure 2B. Bar = 1 µm **(C)** Representative images of the position of the centriole (Asl, Green) relative to the nucleus (DAPI, Red) in RSTs of indicted genotypes. Cartoons (left column) illustrate proper centriole-nuclear docking (≥ 120°, top) vs. an incorrect docking (<120°, bottom). Angle measurement is indicated with black lines. The percent of correct and incorrect docking events for each genotype is indicated (top left corner of each image). The nucleus (n, DAPI, red) is outlined in white. Staining for mitochondria to highlight the derivate/Nebenkern (md) was performed for some samples. FL (n = 64), Δ1 (n = 74), Δ2 (n = 67), Δ12 (n = 116), Δ123 (n = 88). At least 6 testes of each genotype were examined. Bar = 2 µm **(D)** Angle of centriole docking in RSTs in C. Bars are mean ± standard deviation. Statistical comparison to PLP-FL by unpaired t-test with Welch’s correction when appropriate. **** p ≤ 0.0001

We next examined if the N-terminal deletions of PLP affected sperm function by assaying male fertility. Individual males expressing *PLP^Δ2^* showed slight reduction in offspring count, while 65% of *PLP^Δ12^* and 59% of *PLP^Δ1-3^* males produced no offspring (Figure 4A). When the stronger *tubulin-Gal4* driver was used to express these constructs throughout germline development, 88% of *PLP^Δ12^* and 82% of *PLP^Δ1-3^* males produced no offspring (Figure 4B). Consistent with these results, the seminal vesicles (SVs), where mature sperm are stored, were filled with sperm in FL, PLP^Δ1^ or PLP^Δ2^ expressing males, but empty in PLP^Δ12^ or PLP^Δ1-3^ males (Figure 4C). Instead, many sperm were found stranded outside the SV, suggesting that even when made, sperm in the PLP^Δ12^ or PLP^Δ1-3^ background are immotile. We note that the defect in centriole docking only partially explains the male sterility in PLP^Δ12^ and PLP^Δ1-3^, as PLP^Δ1^ and PLP^Δ2^ have similar docking defects (Figure 3D) but are only slightly subfertile. It is possible that the extremely high cytoplasmic levels seen in PLP^Δ12^ and PLP^Δ1-3^ have additional effects on sperm formation, motility, or function; details of which are yet to be explored.

**Figure 4.**
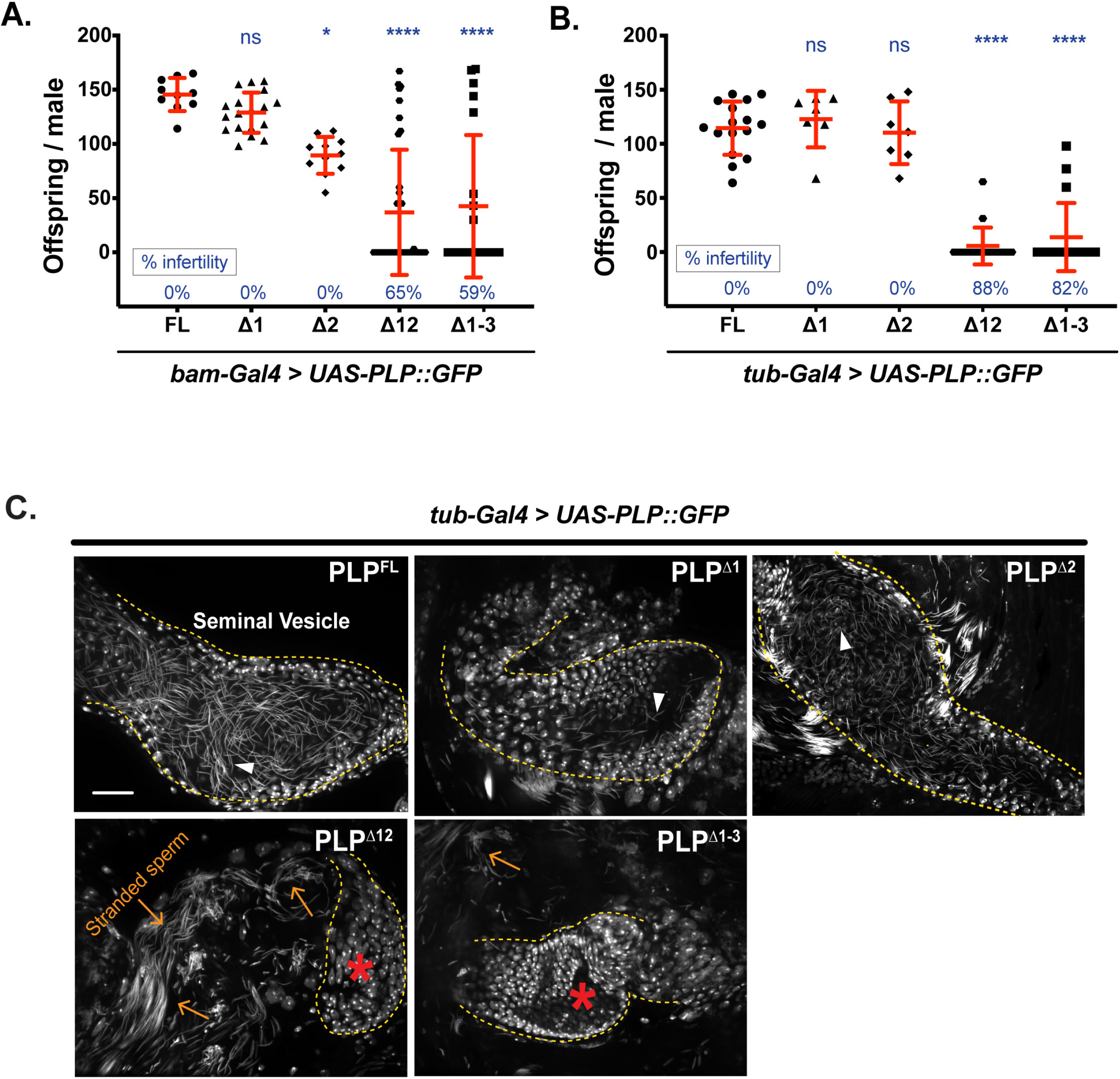
Expression of N-terminal deletions of PLP::GFP impairs fertility. **(A)** Adult progeny per single male mating assay using *bam-Gal4* > *UAS-PLP::GFP* males. Percentage of infertile individuals are indicated along X-axis. Bars represent the mean ± standard deviation. Number of individual crosses for FL (n = 10), Δ1 (n = 18), Δ2 (n = 11), Δ12 (n = 37), Δ123 (n = 21). Statistical comparison to *bam-Gal4>UAS-PLP-FL* by 2-way ANOVA with Dunnett’s multiple comparisons test is shown with * p = 0.0142, **** p ≤ 0.0001, ns = not significant. **(B)** Adult progeny per single male mating assay using *tubulin-GAL4* > *UAS-PLP::GFP* males. Percentage of infertile individuals are indicated along x-axis. Bars represent the mean ± standard deviation. Number of individual crosses for FL (n = 16), Δ1 (n = 7), Δ2 (n = 7), Δ12 (n = 17), Δ123 (n = 17). Statistical comparison to *tubulin-Gal4>UAS-PLP-FL* by 2-way ANOVA with Dunnett’s multiple comparisons test was shown with **** p ≤ 0.0001, ns = not significant. **(C)** Seminal vesicles (SV) from *tubulin-GAL4* > *UAS-PLP::GFP* males stained with DAPI. SV of testes expressing FL, Δ1 or Δ2 were filled with mature sperm (arrowheads). SV of Δ12 or Δ123 contained no sperm (Asterisks), instead the matured sperm were found stranded (orange arrows) outside the SV. N = 5-8 SVs per genotype. Bar =30 µm

### PLP levels are in part controlled by proteosomal degradation

Our results thus far indicated that the N-terminus (PLP^F1+F2^ aa 1-1376) is required to regulate PLP levels in the cytoplasm and on the centriole, and this misregulation can compromise spermatogenesis. To investigate the mechanism of post-translational control of PLP, we turned to cultured S2 cells to perform drug treatments. We first tested PLP turnover by treating S2 cells with cycloheximide (CHX) to block new protein synthesis. Six hours of CHX treatment resulted in near undetectable levels of PLP (Figure 5A, B; Figure S3A), revealing an efficient PLP protein degradation process. We then tested if PLP is degraded via a proteosome-mediated pathway by using the proteosome inhibitor MG132 and found a slight increase in PLP levels (Figure 5A, B, Figure S3B), suggesting the proteosome is required to restrain PLP levels. Finally, we found that treatment of cells with MG132, followed by CHX, resulted in stabilization of PLP protein (Figure 5A, B), indicating a major role of the proteosome in PLP degradation.

**Figure 5.**
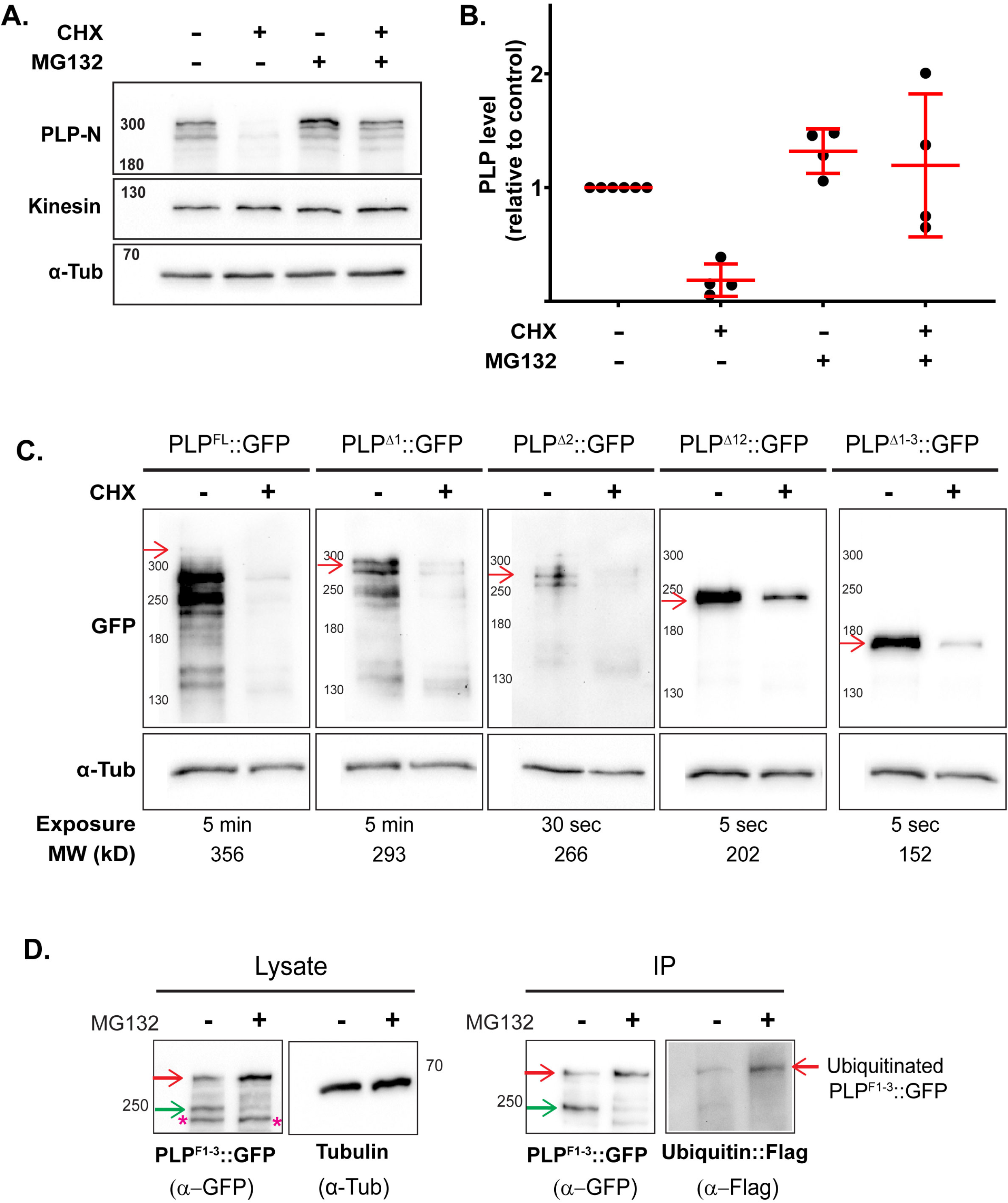
PLP is targeted for proteosomal degradation via sequences in its N-terminal region. **(A)** Western blot of S2 cell extracts showing endogenous PLP level under the following drug treatment condition: DMSO - 6 hours (lane1), Cycloheximide - 6 hours (lane 2), MG132 - 6 hours (lane3), and dually treated – 2 hours MG132 followed by 4 hours with MG132 and Cycloheximide (lane 4). Kinesin and alpha-tubulin are loading controls. The blots were repeated four times. **(B)** Quantitative analysis of PLP level from blots in (A). Signals were normalized to kinesin loading control. Values are relative to PLP level in DMSO control. **(C)** Western blots of S2 cell expressing the indicated PLP::GFP transgenic proteins following GFP pulldown. Cells were treated with DMSO or CHX for 6 hours. Anti-alpha-tubulin was used to confirm input lysates were of similar concentration. Arrow highlight position of PLP band. Expected transgene sizes and blot exposure times are indicated below. Western blots sizes are kDa. **(D)** Western blot of S2 cell immunoprecipitated samples showing the ubiquitination of PLP^F1-3^ after DMSO or MG132 treatment. GFP tagged N-terminal region of PLP^F1-3^ was co-expressed with 3xFlag tagged Ubiquitin. PLP^F1-3^::GFP was IP-ed with anti-GFP and PLP Ubiquitination was identified by blotting for FLAG. Lysate (pre-IP) was blotted for GFP to assess the total PLP^F1-3^ level. Anti-alpha-tubulin of the input confirms similar amounts of lysate were used. Molecular weights are in kDa. We detected three main bands – one at the predicted size of PLP^F1-3^, 250 kDa (green arrow); a lower molecular weight band of unknown origin at 200 kDa (pink asterisks); and a higher molecular weight ubiquitined PLP^F1-3^ at 300kDa band (red arrow).

To determine which regions of PLP contain degradation motifs, we repeated the cycloheximide treatment in cells expressing PLP deletion constructs. Exogenously expressed PLP^FL^::GFP was detected in low abundance, likely a result of the tight control of PLP levels. Due to its low abundance we immunoprecipitated PLP prior to Western blotting. Like the endogenous protein, exogenously expressed PLP^FL^ disappeared after 6 hours of treatment, indicating its rapid degradation (Figure 5C, blot 1). We confirmed this result by Western blot of whole cell extract (Figure S3C). In contrast to PLP^FL^, each of the PLP deletions showed some level of PLP remaining after cycloheximide treatment, indicating that loss of each of these regions reduced PLP degradation (Figure 5C, blots 2-5). This slower degradation rate explains the increase in steady-state concentration of PLP in testes (Figure 2E) and S2 cells (Figure S3D). Together these data indicate that no single region of PLP is sufficient to regulate its level in cells, but instead degradation-regulation is dispersed throughout PLP.

Having noticed that the N-terminal half (aa 1-1811) of PLP is required for its degradation, we next examined if the N-terminal region is ubiquitinated. We transfected S2 cells with PLP^F1-3^::GFP plus 3x Flag-tagged Ubiquitin (Flag-Ubi). These cells were then split into two populations, one was treated with DMSO as a control while the other was treated with MG132. Interestingly, we observed three main bands – one at the predicted size of PLP^F1-3^, 250 kDa; a lower molecular weight band of unknown origin at 200 kDa; and a higher molecular weight band at 300kDa band, which we hypothesized to be post-translationally modified PLP^F1-3^. As predicted, treatment with MG132 elevated the levels of PLP^F1-3^, but in a striking way. The band at 300kDa accumulated while the 250kDa band was no longer detected, consistent with the hypothesis that in the absence of degradation, all of PLP^F1-3^ at 250kDa was post-translationally modified resulting in accumulation of the 300kDa product. To test if this higher molecular weight band was ubiquitinated PLP^F1-3^, we immunoprecipitated (IP) PLP^F1-3^::GFP and blotted for flag-Ubi. This blot showed that the higher molecular weight form indeed represented ubiquitinated PLP^F1-3^. These data support the model that the N-terminus of PLP is ubiquitinated and targeted for degradation via the proteosome.

### Multiple proteasomal components contribute to PLP degradation

To elucidate the mechanisms of PLP degradation, we performed a candidate RNAi knockdown screen for components of the ubiquitin proteasome pathway. We hypothesized that loss of a protein required for PLP degradation would elevate PLP levels in spermatocytes (SCs) during centriole elongation, resulting in precocious PLP localization beyond the proximal end of the centriole toward the distal end (Figure 6A). The screen of 73 candidates included components of the 19s/20s proteosome, ubiquitin E2 conjugating enzymes, substrate targeting E3 ligases previously linked to centrosome biology, and other components associated with protein ubiquitination (Table 1 and 2). Our screen also included eight E3 ligases we identified as possible PLP protein binding partners via an IP of PLP^F2+F3^ (aa 583-1810) followed by mass spectrometry (Table 3, Methods). All UAS-RNAi constructs were driven in early spermatogenesis using *bam-Gal4*; endogenous PLP was evaluated using a C-terminal mNeon CRISPR knock-in (Galletta, 2020). We then used light microscopy to identify gene knockdowns that resulted in distal extension of PLP on centrioles.

**Figure 6.**
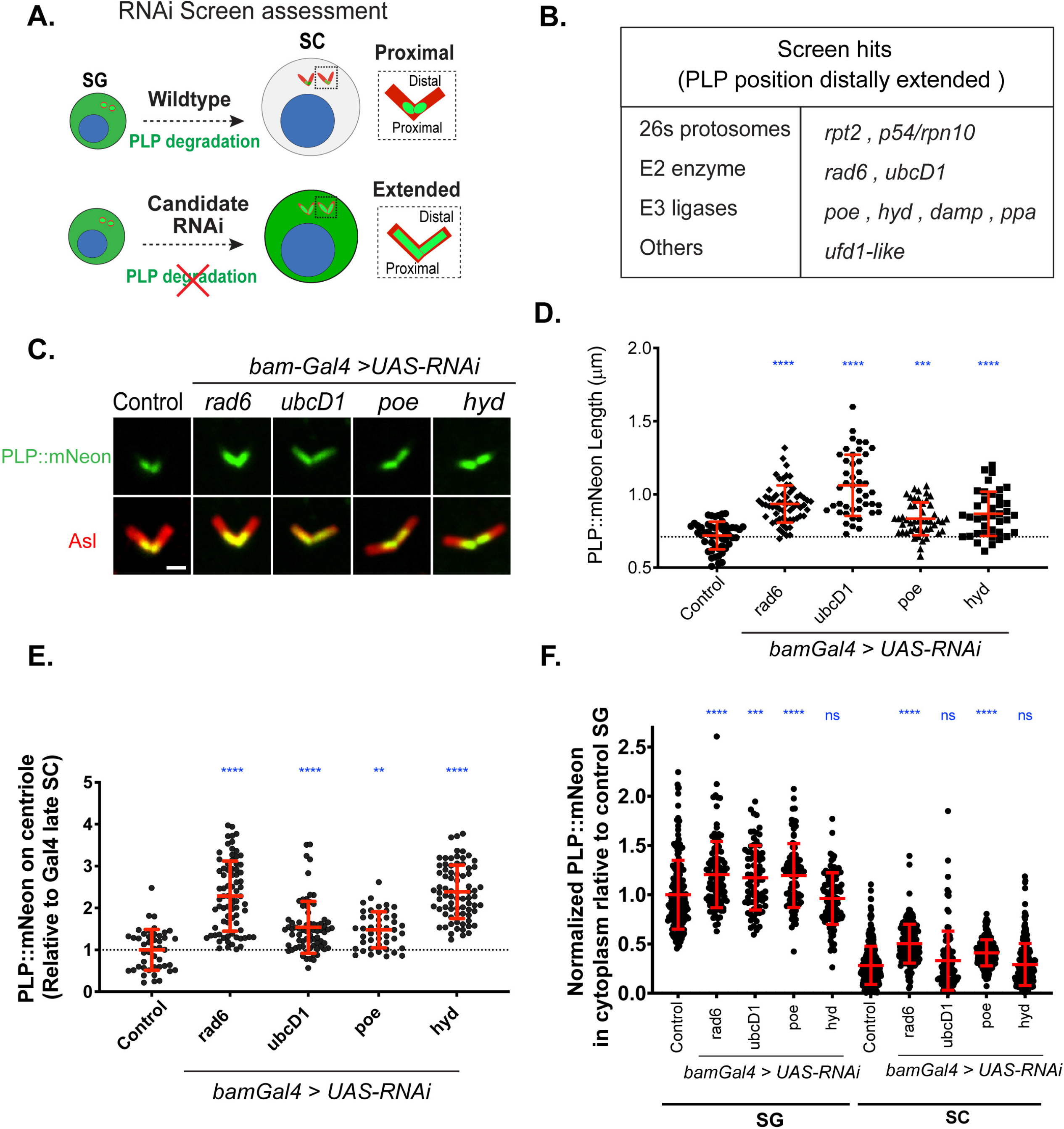
Identification of multiple components that regulate PLP level. **(A)** Cartoon of RNAi screen rationale. Wildtype (top) PLP (green) is actively degraded leading to its proximal localization on the centriole (red). Upon depletion of a hypothetical regulator of PLP degradation (bottom), PLP cytoplasmic levels increase leading to an extension of its position to distal portions of the centriole. **(B)** Genes identified in the RNAi screen to regulate PLP proximal position in spermatocytes. The candidates are grouped into four categories based on function. **(C)** Position of PLP::mNeon (endogenous tag, green) on centrioles (Asl, red) in late spermatocytes from flies expressing RNAi, under control of *bam-Gal4,* against the following genes: Gal4 alone (control), Rad6, UbcD1, Poe or Hyd. Note the extension of PLP on centrioles. Bar = 1 µm **(D)** Measurement of PLP::mNeon length on late spermatocyte centrioles for the *bam-Gal4* control and the indicated knockdowns. Each genotype was examined using 8-10 testes from 4 to 6 flies. Bar represents the mean ± standard deviation. Control mean length is indicated (dotted line). Statistical comparison to control by ordinary one-way ANOVA with Dunnett’s multiple comparison tests with ****p≤0.0001, ***p=0.0003 **(E)** Fluorescent intensity of endogenous PLP::mNeon on centriole from RNAi depleted late spermatocytes for the *bam-Gal4* control and the indicated knockdowns. The measurements are relative to the control level. Each genotype was examined using 8-10 testes from 4 to 6 flies. Bar represents mean ± standard deviation. Control mean length is indicated (dotted line). Statistical comparison to control by ordinary one-way ANOVA with Dunnett’s multiple comparison tests with ****p≤0.0001, **p=0.0019 **(F)** Cytoplasmic level of endogenous PLP::mNeon measured from spermatogonia and late spermatocytes. The individual measurements are shown relative to the cytoplasmic level PLP::mNeon in spermatogonia of control. Bar represents mean ± standard deviation. Statistical comparison by t-test with Welch’s correction when appropriate. ****p≤0.0001, ***p≤0.001, ns = not significant.

**Table 1:**
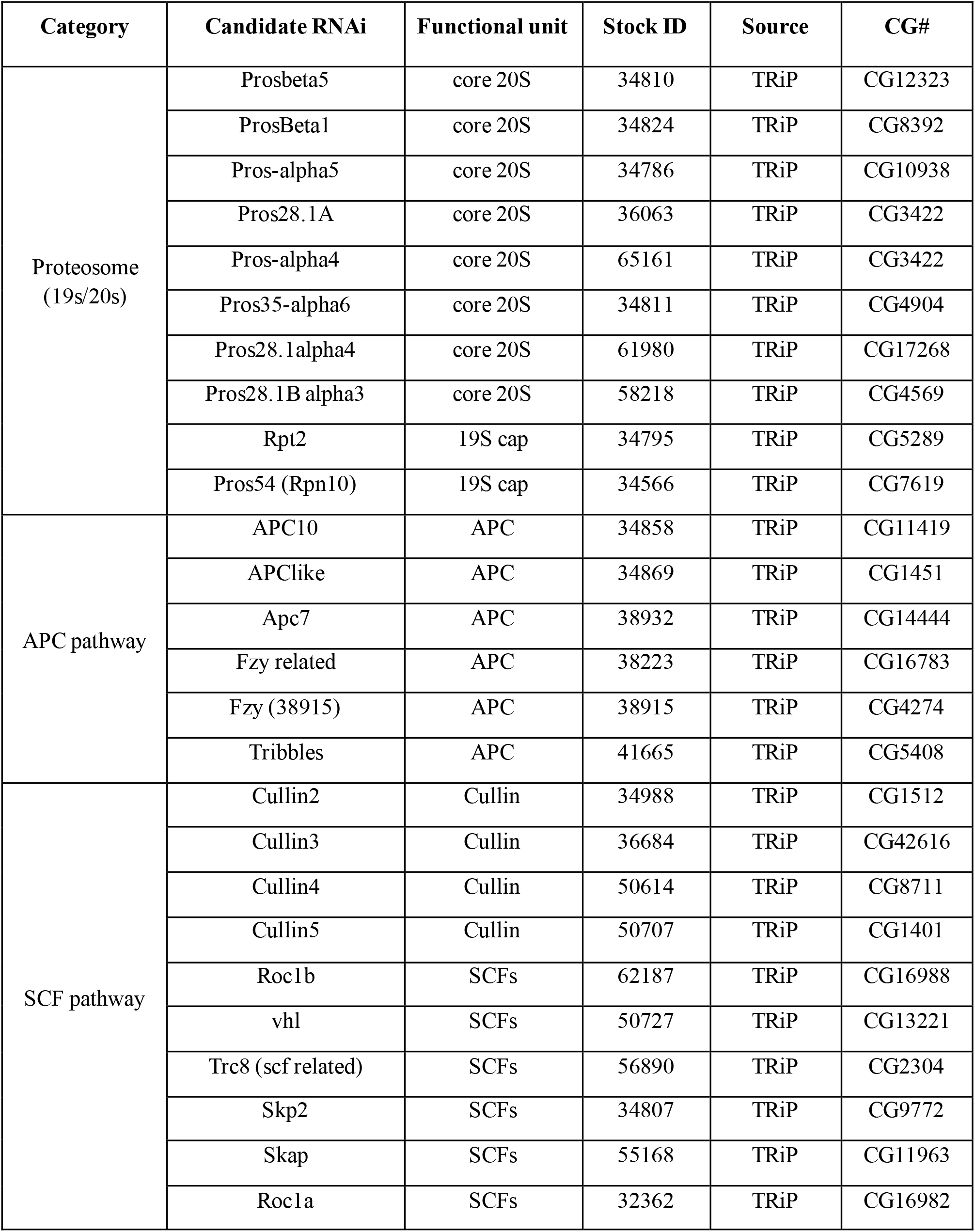
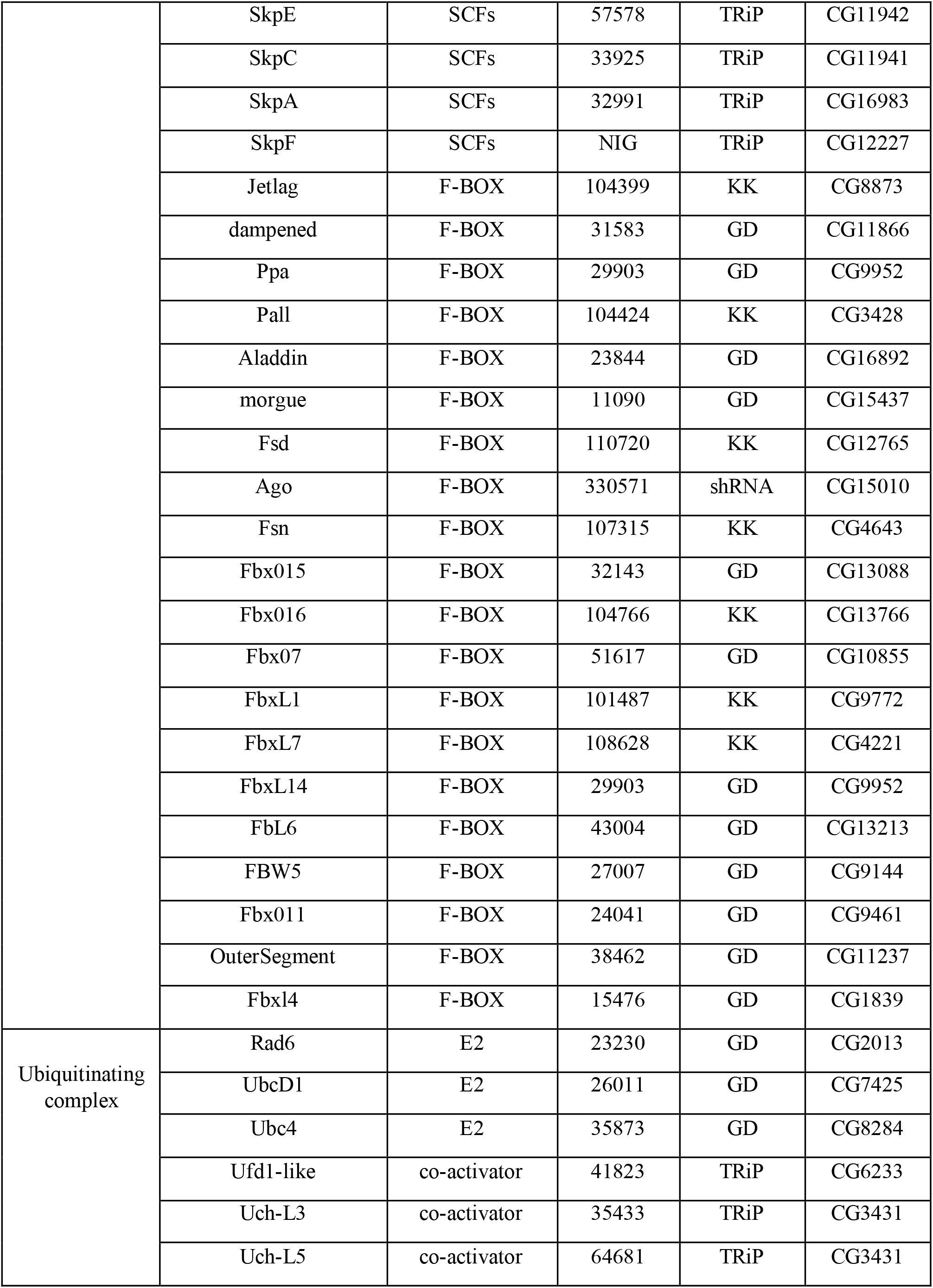
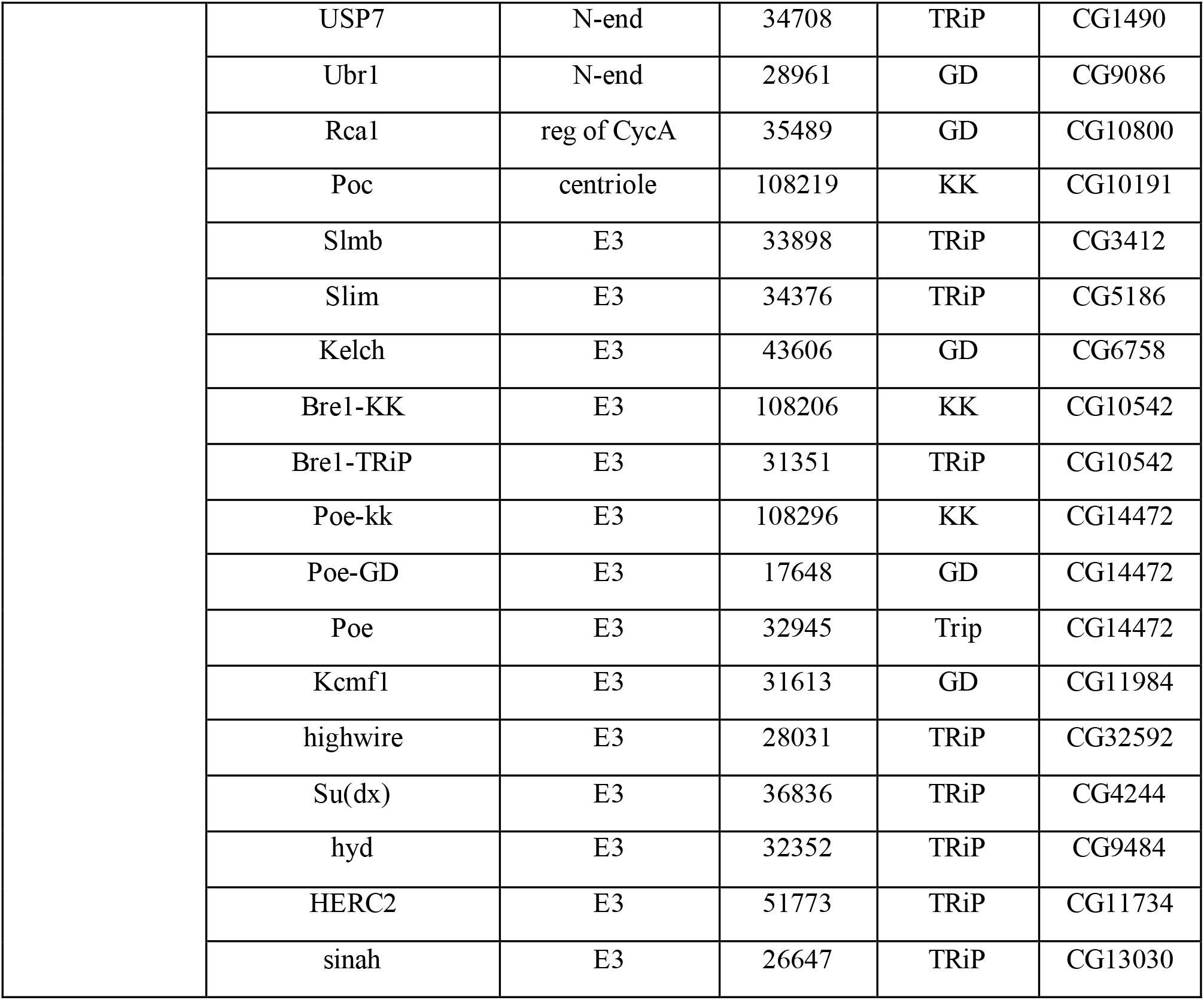
List of candidate RNAi lines tested for PLP position in late SC.

**Table 3:** IP-MS showing PLP interactors that are related to degradation machinery.

| **Description** | **% Coverage** | **# Peptides** | **Score** |
| --- | --- | --- | --- |
| PLPF2F3 | 89 | 170 | 9554.79 |
| E3 ubiquitin-protein ligase highwire | 1 | 1 | 8.76 |
| E3 ubiquitin-protein ligase Bre1 | 2 | 1 | 2.31 |
| E3 ubiquitin-protein ligase Ubr3 | 1 | 1 | 0 |
| E3 ubiquitin-protein ligase Su(dx) | 3 | 1 | 2.67 |
| Protein purity of essence | 3 | 6 | 11.24 |
| E3 ubiquitin-protein ligase hyd | 1 | 1 | 2.31 |
| Probable E3 ubiquitin-protein ligase HERC2 | 1 | 1 | 2.89 |
| Probable E3 ubiquitin-protein ligase sinah | 7 | 1 | 5.57 |
| Cullin-associated NEDD8-dissociated protein 1 | 5 | 2 | 6.8 |
| Frizzled-3 OS=Drosophila melanogaster | 5 | 1 | 2.9 |
| Ubiquitin-like modifier-activating enzyme 5 | 4 | 1 | 2.37 |
| Proteasome-associated protein ECM29 homolog | 2 | 2 | 8.48 |
| F-box/WD repeat-containing protein 7 | 2 | 1 | 5.87 |
| Probable ubiquitin carboxyl-terminal hydrolase | 1 | 1 | 6.48 |

Our qualitative screen for extended PLP positioning yielded 9 hits (Figure 6B); additional gene knockdowns resulted in other phenotypes such as centriole length abnormalities or formation of PLP cytoplasmic aggregates (Table 2, not further studied here). Of the 9 hits, we selected 4 genes for additional quantitative analysis: two E2 conjugating enzymes (Rad6^(E2)^ and UbcD1^(E2)^) and two E3 ubiquitin ligases (Poe^(E3)^ and Hyd^(E3)^). These genes were selected because they have been previously shown to regulate protein levels via N-terminal regions (see discussion). Follow-up protein knockdown of Rad6^(E2)^, UbcD1^(E2)^, Poe^(E3)^, or Hyd^(E3)^ resulted in increased PLP::mNeon length along the centriole (Figure 6C, D), confirming the initial screen result. Additionally, all four knockdowns increased PLP levels on centrioles (Figure 6E). Quantitative fluorescence analysis of the cytoplasm showed that knockdown of Rad6^(E2)^, UbcD1^(E2)^ and Poe^(E3)^ lead to elevation of PLP levels in the cytoplasm of SG, while knockdown of Rad6^(E2)^ and Poe^(E3)^ maintained high PLP in the cytoplasm of SC (Figure 6F). These results suggest that, to different extents, PLP degradation is regulated by Rad6^(E2)^, UbcD1^(E2)^, Poe^(E3)^, and Hyd^(E3)^, possibly directly targeting distinct PLP regions.

**Table 2:**
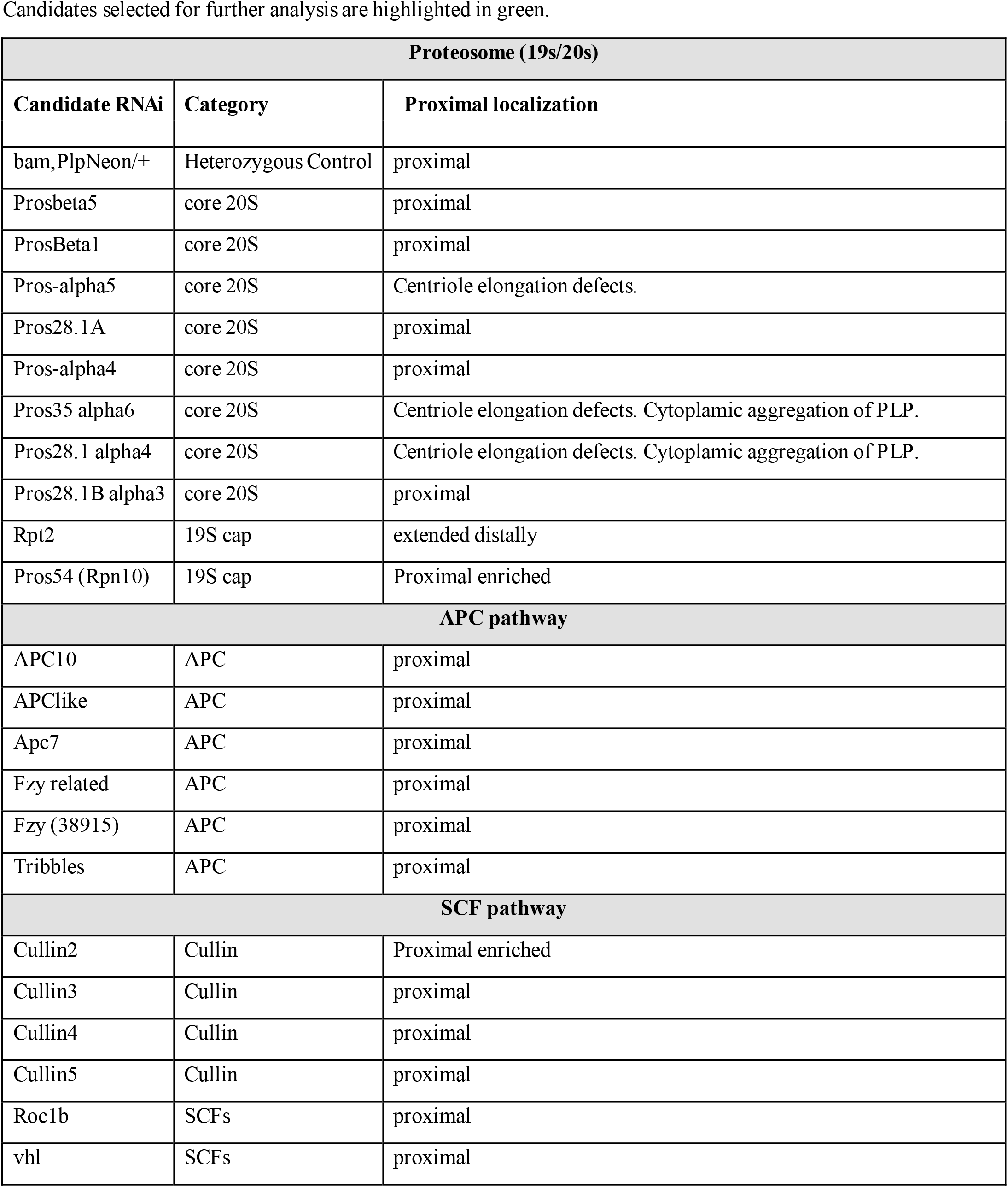
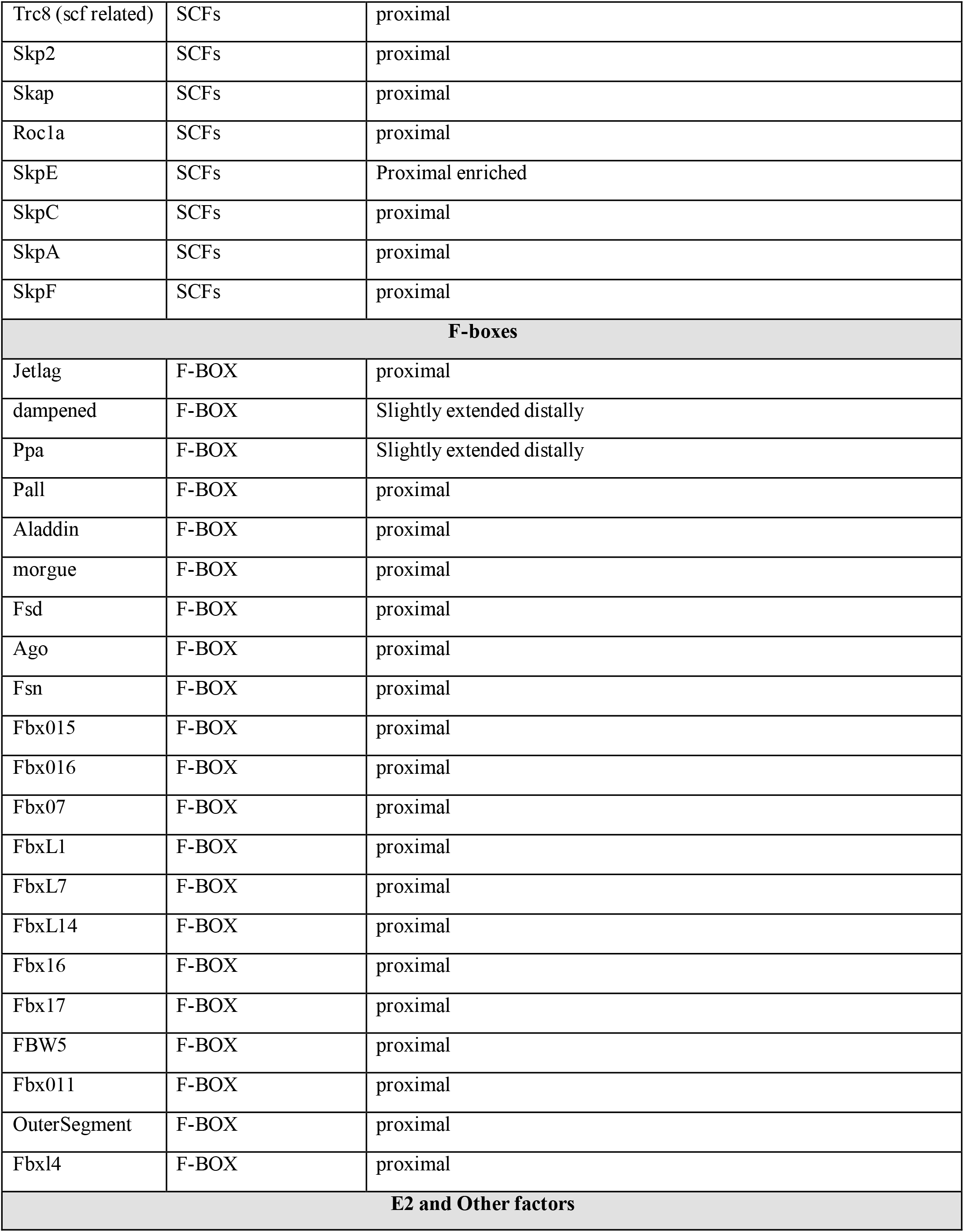
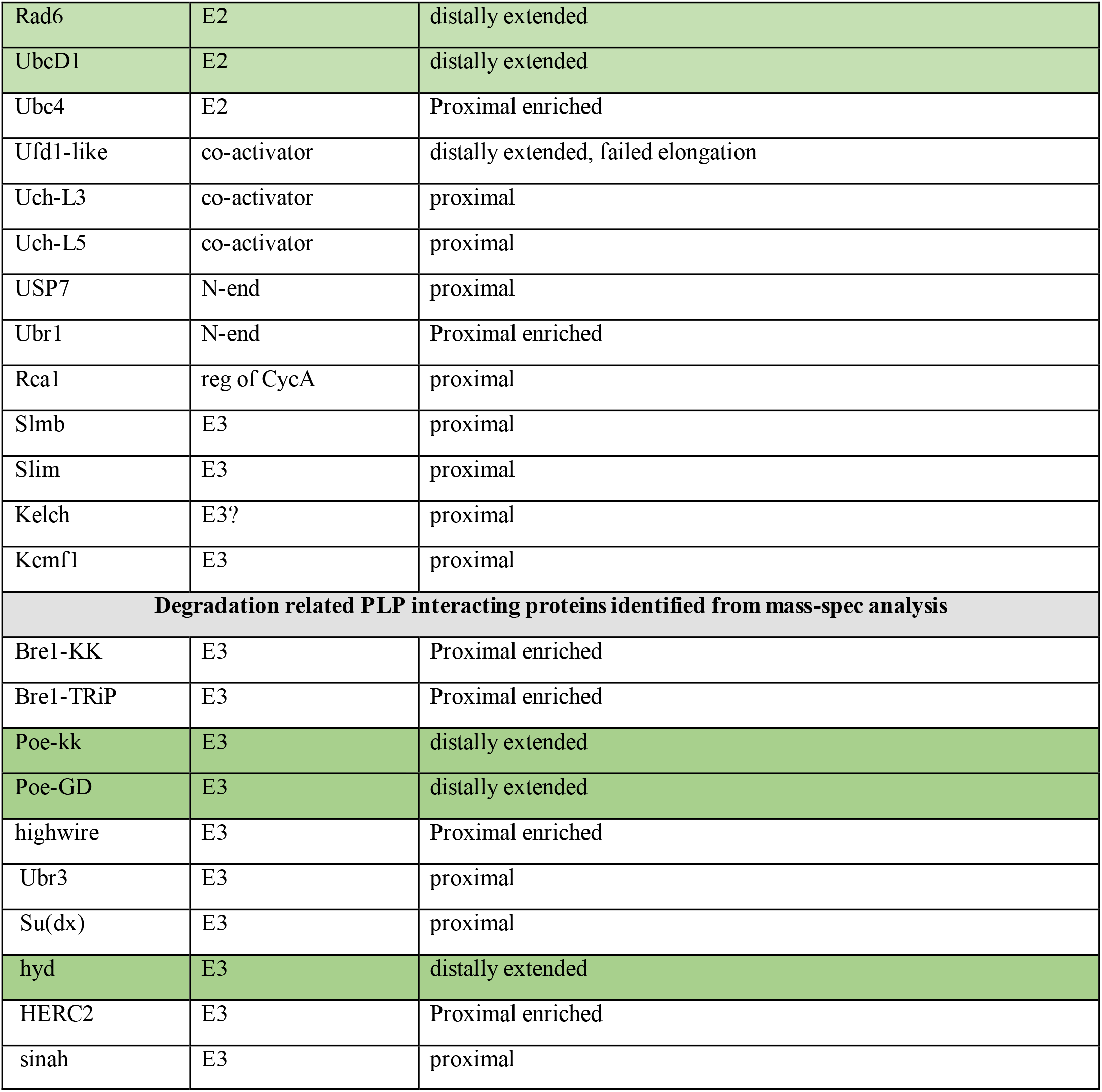
Summary of Candidate RNAi screen for PLP position in late SC.

### PLP directly binds Poe and Hyd

To test if PLP is a direct substrate of Poe and Hyd, we used two complementary assays to identify possible protein-protein interaction – protein immunoprecipitations (IPs; Figure 7 A, C) and Y2H (Figure 7 B, D). We performed immunoprecipitations from cells co-expressing PLP^F1-3^::GFP fragment and Flag tagged Poe^(E3)^ or Hyd^(E3)^ fragments in S2 cells. We tested the interactions using fragments of Poe^(E3)^ covering its entire length and fragments of Hyd^(E3)^ containing conserved and previously predicted functional domains (Cammarata-Mouchtouris, 2020). We found that the N-terminal region of PLP (aa 1-1811) co-IPed with all tested regions of Poe, including the region containing the Zn finger (ZnF, aa1621-2040) domain located in fragment 2 of Poe (Figure 7A, E), which can serve as a type 1 substrate recognition domain of N-recognins (Tasaki, 2007; Tasaki, 2012). Our Y2H assay showed that PLP^F2-F3^ (aa 584-1811) interacted with Poe^F1^ (aa 1-1100; Figure 7B) and Poe^F2^-ZnF (aa 1621-2040; Figure 7B). The differences seen between the IP data and the Y2H interactions in Poe ^F3^ and Poe^F4^ might suggest indirect protein interactions, the reliance on a missing post-translational modification, or simply an uninterpretable Y2H negative result. Additional Y2H experiments using the C-terminal end of PLP^F5^ (aa 2539-2895) revealed direct interaction with the conserved ZnF of Poe^F2^ (aa 1741-1920; Figure S4, row 6), but we did not pursue this interaction further.

**Figure 7:**
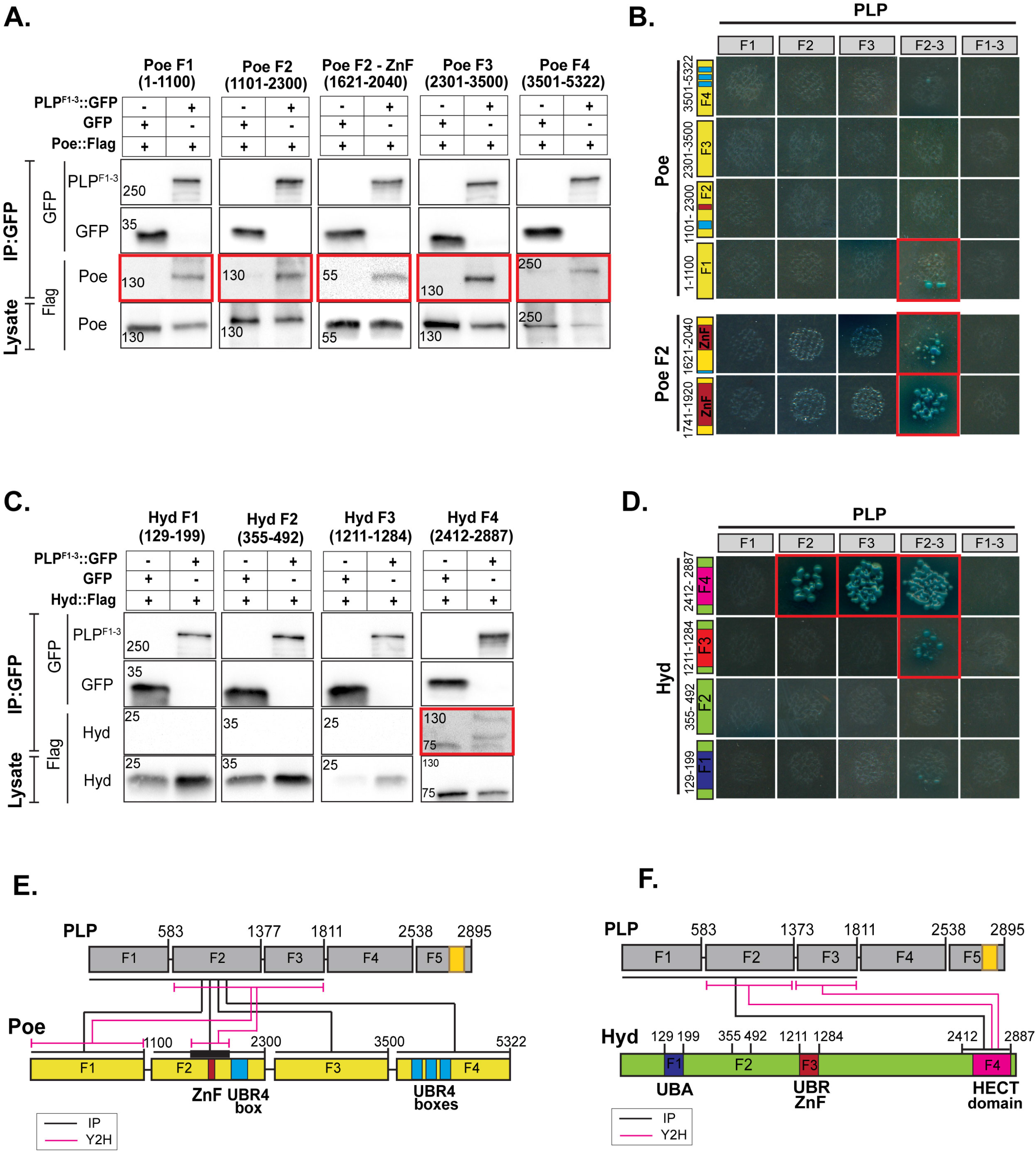
Interactions between PLP and the E3 ligases Poe and Hyd. **(A)** GFP tagged N-terminal region of PLP (F1-3) was co-expressed with Flag tagged Poe-F1 (aa 1-1100), Poe-F2 (aa 1101-2300), Poe-F2-ZnF (aa 1621-2040), Poe-F3 (aa 2301-3500) or Poe-F4 (aa 3501-5322) in S2 cells. PLP was IP-ed with anti-GFP and blotted for GFP and FLAG. Pulldown of GFP was used as a control. Poe::Flag fragments co-IP-ed with PLP (F1-3) are indicated in red boxes. Lysates (pre-IP) show Poe::Flag expression in both GFP and PLP (F1 -3) samples. **(B)** Yeast two hybrid (Y2H) interaction assay between the N-terminal fragments of PLP (F1 to F3) and fragments of Poe-F1, F2, F2-ZnF, F3 or F4. Amino acids of Poe included in each fragment indicated in figure. Growth and blue color indicate interaction (red boxes). **(C)** GFP tagged N-terminal PLP (F1-3) was co-expressed with Flag tagged fragments of Hyd F1(aa 129-199), F2(aa 355-492), F3(aa 1211-1284) or F4 (aa 2412-2887) in S2 cells. PLP was IP-ed with anti-GFP and blotted for GFP and Flag. Pulldown of GFP alone was used as a control. Hyd F4 co -IPed with PLP F1-3 (red box). Lysates (pre-IP) show Hyd::Flag expression in all samples. **(D)** Y2H interaction assay between the N terminal fragments of PLP (F1 -3) and fragments of Hyd-F1, F2, F3 or F4. Growth and blue color indicate interaction (red boxes). **(E)** Schematic of the Y2H (pink lines) and IP (black lines) interactions between PLP and Poe. **(F)** Schematic of the Y2H (pink lines) and IP (black lines) interactions between PLP and Hyd.

We performed similar IP and Y2H assays to investigate a possible PLP and Hyd interaction. Both IP and Y2H revealed that the C-terminal catalytic domain of Hyd, containing the Hect substrate recognition domain (aa 2484-2885), bound directly to each of PLP^F2^ and PLP^F3^ (Figure 7C, D, F). These interactions support a model that PLP is a substrate of Poe^(E3)^ and Hyd^(E3)^.

## Discussion

A critical aspect of the multi-faceted process by which cells control protein levels, and in turn cellular homeostasis, is protein degradation. Cells not only use protein degradation to maintain proteins at specific, optimal concentrations, they also use protein degradation to regulate dynamic processes in time and space, including the progression of the cell cycle, cell signaling, metabolism and many others (Walter, 2011; Yoo, 2018). For example, protein degradation can fine tune the timing of a cellular response, ensuring that it is muted or active at the appropriate time. Defects in protein degradation can perturb cellular homeostasis and result in disrupted cellular functions that have been linked to a development of a variety of disease conditions like Alzheimer’s, neuronal degenerative disease, cancer, etc. (Ciechanover, 2015; Hipp, 2019; Nakayama, 2006; Watanabe, 2020).

Centrosomes have long been recognized as hubs where protein degradation components are localized, including the APC and SCF (Freed, 1999; Torres, 2010; Tugendreich, 1995). However, it also appears that the function of the centrosome itself is regulated by specific and timely degradation of its regulators and component proteins (see introduction for more detail). This is not surprising given the tight coordination between the cell cycle and each of centrosome duplication (S-phase), centrosome activation (late G2), and centrosome inactivation (early G1). For example, core centrosome regulators and component proteins such as Plk4 (Guderian, 2010; Klebba, 2015; Klebba, 2013; Rogers, 2009), Sas6 (Puklowski, 2011; Strnad, 2007), and Polo (Lindon, 2004) are all subject to regulated degradation to ensure that they are only present when needed. It has also become apparent that the titer of centrosome proteins and their specific modification is critical for both the assembly and disassembly of centrosomes (Alvarez-Rodrigo, 2019; Woodruff, 2017; Woodruff, 2015), although the role of protein degradation in these process has yet to be examined.

One protein whose levels must be regulated to ensure proper function is Pericentrin (PCNT in humans and PLP in *Drosophila*). PCNT was found to be hyper accumulated in tumor cells from pancreatic and prostate cancer patients and it was suggested to be the cause for the characteristic ectopic MTOC formation and MT nucleation in these cells (Kim, 2008; Sato, 1999). Elevation in PCNT expression has been reported in patients affected with Down syndrome and bipolar disorders (Anitha, 2009; Salemi, 2013). Cells harboring trisomy 21 have abnormal PCNT accumulations and defects in cilia formation and function (Galati, 2018). Consistent with these observations, artificial elevation of PCNT levels results in defects in MTOCs, abnormal spindles and aneuploidy (Purohit, 1999). Together, all these studies indicate the relevance and the requirement for regulating PCNT level, however, the mechanisms by which the PCNT level is regulated remains unclear. Interestingly, expression of portions of PCNT or the PCNT related protein AKAP350 (PCNT related) form cytoplasmic aggregates, which in turn recruit PCM components and nucleate microtubules independent of centrosomes. These studies suggest that regions of PCNT/AKAP350 are required to limit PCM assembly to the proper place and time, however the regulatory mechanisms that impose these limits remain unexplored (Varadarajan, 2017; Kolobova, 2017; Jiang, 2021).

We had previously demonstrated that PLP levels are sharply downregulated during early spermatogenesis and that this regulation is essential to spatially position PLP at the proximal end of the long centrioles in *Drosophila* SCs (Galletta, 2020). While we showed that the mRNA encoding PLP was only available in early germ cells, the mechanism by which protein levels dropped remained unclear. Here we demonstrate that PLP is subject to proteasomal degradation that acts alongside the previously identified mRNA regulation to temporally coordinate the clearance of PLP in developing SCs. Both these mechanisms appear to directly influence the proximal localization of PLP and therefore the spatial positioning of PCM on centrioles, which in turn influences proper centriole docking in spermatids (Figure 8).

**Figure 8:**
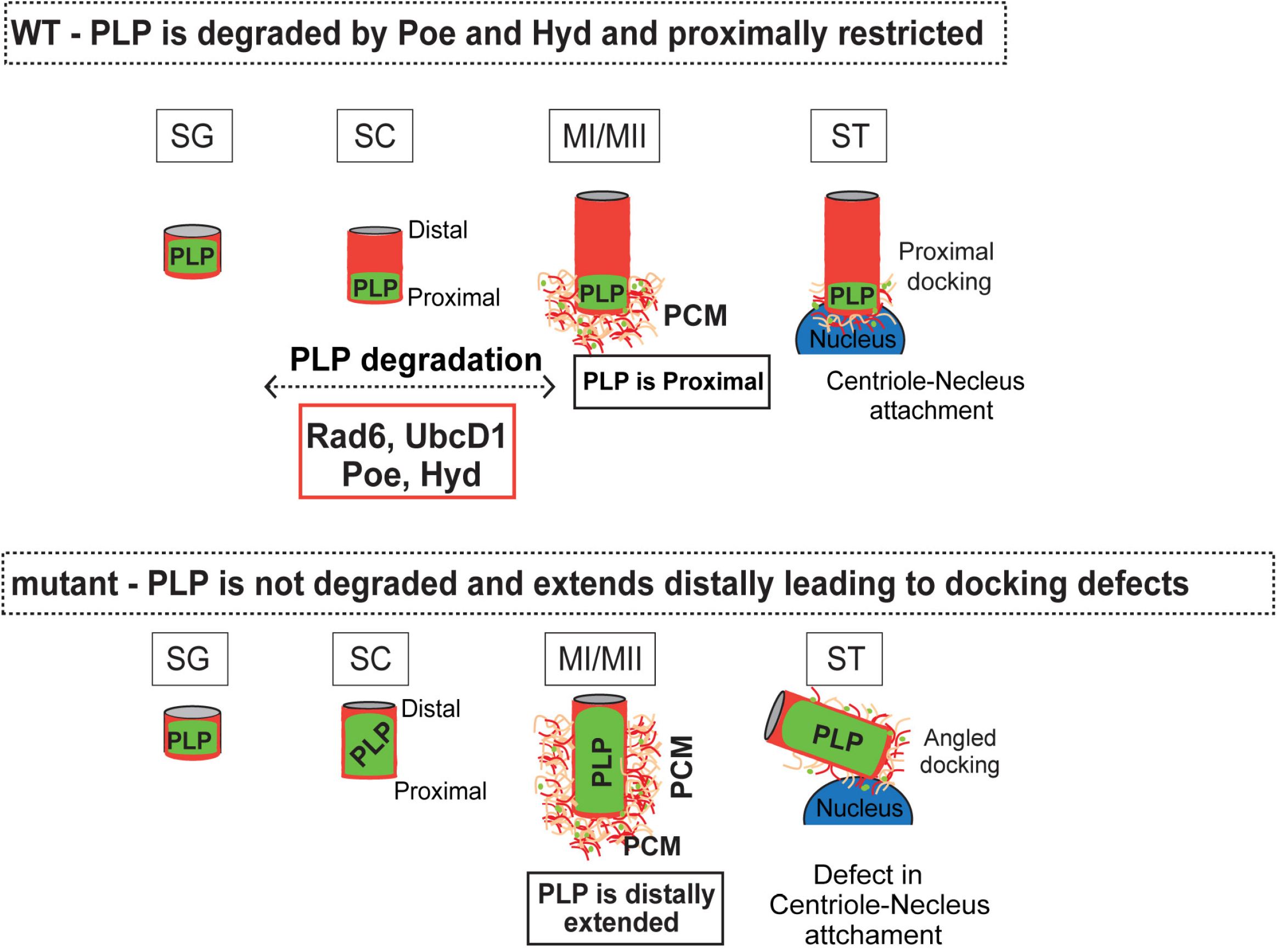
Summary Model of PLP degradation by Poe and Hyd (WT) Normally cells efficiently degrade PLP, in part via Poe and Hyd, prior to centriole elongation. **(Mutant)** Disruption of Rad6, UbcD1, Poe, Hyd prevents efficient PLP degradation, resulting in defective centriole-nucleus docking. This

Our data suggests that post-translational regulation of PLP is complex. Multiple regions of the protein appear to contain sequences that regulate its degradation and multiple ubiquitin ligase components appear to be involved. Moreover, our screen identified Rad6^(E2)^ (Dohmen, 1991), UbcD1^(E2)^ (Treier, 1992), Poe^(E3)^ (Ashton-Beaucage, 2016; Kim, 2018) and Hyd^(E3)^ (Cammarata-Mouchtouris, 2020; Dorogova, 2020; Kinsella, 2016) as regulators of PLP. Poe^(E3)^ (human UBR4) and Hyd^(E3)^ (human UBR5) are N-recognins, members of a family of UBR box containing E3 ligases known to recognize the substrates of N-end rule ubiquitination system (Kim, 2018; Tasaki, 2009). Interestingly, Rad6^(E2)^ is known to function with Poe^(E3)^ in its N-end rule regulation of proteins involved in vesicular trafficking and the lysosomal pathway (Hong, 2015). Hyd^(E3)^ has been studied in context of the N-end rule in regulating Hedgehog signaling, oogenesis, and immune response in *Drosophila* (Cammarata-Mouchtouris, 2020; Dorogova, 2020; Moncrieff, 2015), while its human ortholog UBR5 was studied in ovarian cancer signaling, tissue homeostasis, and miRNA mediated translational control (Cho, 2017; Hughes, 2021; Mellis, 2021; Tasaki, 2012). Finally, while UbcD1^(E2)^ has not been linked to the N-end rule, it does show high similarity to UBC4 (Treier, 1992), which is an E2 known to function with Hyd^(E3)^/UBR5 in yeast (Stoll, 2011). Thus, we hypothesize that PLP could be regulated by the N-end rule via a degron recognized by Rad6^(E2)^-Poe^(E3)^ and/or UbcD1^(E2)^-Hyd^(E3)^.

Typically N-end rule substrates require single or sequential enzymatic modifications of the very N-terminal residues to form N-degron (Varshavsky, 2017). However, the PLP interactions we identified with Poe and Hyd suggest the presence of internal degrons in PLP. Studies from yeast have shown that the UBR recognins of Arg/N-end rule pathway not only recognize the N-degrons of their substrates, but also internal degrons (Varshavsky, 2019; Xia, 2008). Given the structural and functional similarity between PLP and PCNT, and the documented importance of controlling its protein levels, we hypothesize that the human homolog of these N-recognins (Poe/UBR4 and Hyd/UBR5) also regulate PCNT degradation. Our future work will focus on testing the N-end rule hypothesis directly.

While we demonstrate that PLP is subject to post-translational regulation via protein degradation, it seems likely that this type of regulation might be used for other centrosome proteins or in other cellular contexts. For example, protein degradation might be used in cycling cells to ensure that PLP is only available at a precise time during the cell cycle to load onto centrioles and might be degraded to ensure that it is not available at other times. Furthermore, while our current study focuses on regulating the availability of the protein in the cytoplasm for assembly onto the centriole, protein degradation might very well be used to remove proteins from the centriole itself, such as the case in the process of ‘Centrosome Reduction’ (Khire, 2016; Manandhar, 2005; Schatten, 2015). This process of stripping proteins like PLP (Galletta, 2020) and Asl (Khire, 2016) from centrioles is likely subject to tight degradation regulation, a topic yet to be explored. We predict that the role of the N-end rule pathway in regulating other centrosome proteins, on and off of the centriole, will be a major area of future exploration.

## Acknowledgements

We thank S. Smith, R. Liu and J. Sellers for help with IP-MassSpec analysis, A. Sodeinde for testes dissections, R. O’Neill, and M. Hannaford for active discussions. We thank A. Kelly and T. Akera and their lab members for the critical discussions. We thank J. Ryniawec and G.C Rogers for advising on PLP blots and for reagents. We thank the Blooming Drosophila Stock Center and the Vienna Drosophila Resource Center for flies and the Developmental Studies Hybridoma Bank for antibodies. We thank the NHLBI Light Microscopy, Biophysics, and Proteomics core facilities for technical support. This work was supported by the Division of Intramural Research at the National Institutes of Health/National Heart, Lung, and Blood Institute (1ZIAHL006104 NMR).

## Materials and Methods

### Generation of Transgenic Flies

A full length PLP (PF isoform) cDNA was used to generate all PLP constructs. The five PLP sub fragments were generated previously (Galletta, 2014). Truncations were generated by PCR of the region of interest (deltaF1 [PCR of F2-5], deltaF1-2 [PCR of F3-5], deltaF1-3 [PCR of F4-5]) followed by pENTR/D TOPO cloning. PLP deltaF2 cDNA was generated as previously described (Lerit, 2015). These were then cloned into the full-length cDNA by restriction digest using NotI and XhoI. cDNAs were cloned into P-element destination vector (pPWG; G=GFP) using Gateway cloning system to express under the control of a UAS promoter (Invitrogen) and to include a C-terminal GFP tag. N-terminal GFP tagged PLP was generated using pPGW vector. Transgenic flies were generated using standard P-element transformation (BestGene; Chino Hills, CA).

### Fly stocks

Flies used in this study were cultured in standard cornmeal-agar media at 25°C. PLP::mNeon is a CRISPR knock-in (Galletta, 2020). Transgenic expression of full length PLP (PLP^FL^) transgene was used as an experimental control for all PLP truncations and N-terminal modified PLP lines. bam-Gal4 (Chen, 2003) was used for testes specific expression and tubulin-Gal4 (Lee, 1999) for ubiquitous expression of UAS-driven transgenes. Ubi-PLP::GFP contains the PLP^FL^ coding sequence with a C-terminal GFP-tag under the control of the ubiquitin promoter (Galletta, 2014). RNAi lines were obtained from Harvard TRiP collection and VDRC stock center (Table 1) for the candidate genetic screen. *bam-Gal4*, *tubulin-Gal4* and all the TRiP lines (Table 1) were obtained from Bloomington stock center and other RNAi lines were obtained from Vienna Drosophila Resource Center.

### Plasmids for Drosophila S2 expression

For the interaction assays, we first synthesized the segments of cDNA sequences covering the entire length of Poe and sequences for the conserved domains of Hyd. The cDNA sequence encoding the following Poe fragments F1 (aa 1-1100), F2 (aa 1101-2300), F2-ZnF (aa 1621-2040), F2-ZnF (aa 1741-1920), F3 (aa 2301-3500) and F4 (aa 3501-5322) and, the sequence of Hyd F1(aa 129-199), F2(aa 355-492), F3(aa 1211-1284) and F4 (aa 2412-2887) were synthesized (Twist Bioscience). The individual cDNA sequences were cloned into pENTR Topo vector. Destination reactions were performed to further clone the E3 cDNA segments into destination vectors containing Actin5C or Ubiquitin promoter to express protein fragment with the desired tags (AGW, AWG, UWG, UGW, AFHW; G=GFP, F=Flag (https://emb.carnegiescience.edu/drosophila-gateway-vector-collection) using the Gateway cloning system (Invitrogen). AWG-MCS plasmids were used as GFP control for IP experiments.

### S2 Cell Transfection and Treatment

Drosophila S2 cells (Invitrogen and DGRC) were maintained in SF900 media (Lif e Technologies) supplemented with 1x Antibiotic-Antimycotic (Life Technologies) at 25°C. Cells were transfected using Nucleofection (Lonza) as recommended by the manufacturer (Amaxa). 1 µg of plasmid was preincubated in 100 µl nucleofection reagent (50 mM D-mannitol, 15 mM MgCl2, 5 mM KCl, and 120 mM NaPO4, pH 7.2) for 15 minutes. Plasmid mix was used to resuspend ∼2 - 5 million cells and cells were electroporated using G-030 program in the Lonza Amaxa nucleofector. Transfected cells were then transferred to 2 ml SF900 media and incubated at 25°C for 48-72 hours before use. Drug treatments: Steady state PLP protein level was analyzed by treating S2 cells with 50µM of MG132 or 50µg/mL of Cycloheximide (CHX) for 6 hours. DMSO without drug was used as control. When both drugs were used, cells were incubated in MG132 for two hours followed by CHX and MG132 for 4 hours. For transfected cells expressing PLP::GFP, drug treatments were begun 48 hours after transfection.

### Immunostaining

Testes were dissected from 1-3 day old males in SF-900 media (Gibco) then fixed in 4% formaldehyde diluted in PBS for 20-30 minutes and permeabilized in 1% PBT (triton x100 diluted in PBS) for 15 minute or 9% formaldehyde in PBS for 20 min followed by brief washes in 0.3% PBT. Samples were blocked in 5% normal goat serum diluted in 0.3-1% PBT at room temperature for 30 min to 4 hours and incubated in primary antibodies diluted in 0.3-1% PBT at the concentration mentioned below, overnight at 4°C. Samples were then 3x washed in 0.3-1% PBT, each for 7-10min and incubated in secondary antibodies diluted in 0.3-1% PBT (see below) for 1-2 hours at room temperature. After washing and counterstaining with DAPI, the testes were mounted on coverslip using Vectashield (VectorLabs) or Aquapolymount (Polysciences, Inc). The following are the concentration of primary antibodies used for immunolabeling in this study: rabbit anti-PLP, raised against the N-terminus region, 1:5,000 (Rogers, 2008); Guinea pig anti-Asl, 1:10,000 (Rogers, 2008); rabbit anti-Cnn (Galletta, 2016), 1:10,000: mouse gTub (GTU-88; Sigma Aldrich), 1:500, anti-ATP5A (1:1000, 15H4C4, ab14748; Abcam). Secondary antibodies labeled with Alexa Fluor 488, 568 or 647 conjugations were used in 1: 1000 dilution (Thermo Fisher Scientific). DAPI (1:1000; Thermo Fisher Scientific) was added to secondary antibodies or the second wash after incubation with secondary antibodies.

### Microscopy

Samples were imaged using a Nikon W1 with a spinning disc confocal head (Yokogawa), 405, 488, 561 and 641 nm laser lines and Prime BSI cMOS camera (Photometrics). Unless noted, images shown in this study were imaged using 100x/1.35 NA silcone immersion objective. Images of testes seminal vesicle were imaged with 40X/1.3 NA water immersion objective. Cytoplasmic measurements were made with a 40X/1.3 NA oil immersion objective. The microscope was controlled, and images were acquired using Nikon Elements software (Nikon). All images were analyzed and processed using FIJI (ImageJ; National Institute of Health).

### Quantification and Statistical analysis

All the data analysis and statistics were performed using Excel (Microsoft) and Prism (Graphpad) software. Statistical analysis was performed using Student’s t tests with Welch’s correction when necessary, or One-way ANOVA with Dunnett’s multiple comparison test, when appropriate. Sample sizes are reported in the figure legends. For quantifications the mean ± standard deviation is presented.

PLP distance measurement: PLP distribution was measured using ImageJ software from the images of centrioles from the late-stage spermatocytes. Asterless label (Red) was used as a marker to identify these centrioles. For characterization of initial screen hits PLP::neon distribution was measured by drawing a line along the length of Asl label covering the proximal to distal region of centriole where PLP (Green) is localized.

Centriole protein level measurement: For quantification of levels, we prepared and imaged all samples on the same day. Fixed samples, with stained with anti-Asl to label centrioles and DAPI to label nuclei. Anti-ATP5A staining was done on some samples to aid in staging spermatids. Sum projections of the entire Z volume of the centriole were generated, region of interest (ROI) was drawn around the centriole using the Asterless label for reference, and the total integrated density was measured. The same ROI was then moved away from the centriole and the intensity was measured for the background subtraction. The measurements were normalized to the mean value of the bam-Gal4 control on a given day.

Cytoplasmic protein level: All were measured in live germline cells performed by mounting the testes on 50 mm lummox dish (Sarstedt) in a drop of S2 media, surrounded by Halocarbon oil 700 (Sigma) and covered with a No. 1.5 coverslip. Single confocal planes were acquired. Cytoplasmic protein level was analyzed by measuring the average pixel intensity of PLP::mNeon or PLP::GFP in a ROI drawn in the cytoplasm, avoiding any large aggregates or centrosomes. The same ROI was then moved off the sample to acquire a background signal for subtraction. All the measurements were plotted relative to the cytoplasmic level of bam-Gal4 driving UAS-PLP::GFP in spermatogonia on a given imaging day. Since aggregates were avoided, these measurements may underrepresent the total cytoplasmic levels.

### Immunoprecipitation

For IPs, S2 cells were harvested two days after transfection and resuspended in lysis buffer (50 mM Tris, pH 7.2, 125 mM NaCl, 2 mM DDT, 0.1% Triton X-100, and 0.1 mM PMSF or 50 mM Tris pH 7.4, 150 mM NaCl, 0.5% Triton X-100, 1 mM DTT, 1 mM PMSF, 1 ug/mL Leupeptin, 1 ug/mL Pepstatin). After 10 -15 min on ice, the lysate was cleared (3 mins, 21130xg, 4°C). The supernatant was then incubated with 15-25 µl of Protein-A dyna beads conjugated with His-Lamma GFP binding proteins (GBP) (Patron, 2019; Rothbauer, 2008) for 0.5 - 2 hours at 4°C with mixing. The GFP tagged protein bound to GBP-dynabeads were washed thrice in lysis buffer on ice, each wash for 1 min, eluted by boiling in 1 or 2x SDS-sample buffer (58 mM Tris pH 6.8, 5% glycerol, 1.95% SDS, 1.55 % DTT, $0.05% Bromophenol Blue) for 5 -10 minutes and analyzed by Western Blot. The input lysate and flowthrough were also analyzed.

### Western blots

For blots of testes extracts, 50 adult testes were dissected in S2 cell media from 1 –3-day old male flies and homogenized in 30 uL of IP lysis buffer, followed by addition of SDS-sample buffer. Samples were boiled for 10 min and stored at -80°C until use. For S2 cells, lysate was prepared by harvesting cells after treatment or transfection followed by lysing cells in IP lysis buffer, normalizing the volume according to the cell count to achieve similar cells/ml. A small sample was taken, and total protein was estimated by Lowry assay (Lowry, 1951). SDS sample buffer was then added, and samples were boiled for 5 min. Volume was adjusted prior to loading to ensure the same amount of total protein was loaded into each lane. Samples were run on 6% or 7.5% SDS-polyacrylamide gels. Samples were transferred to nitrocellulose or PVDF membrane using Tris-Base Glycine transfer buffer (Novex) with 20% Methanol. Blots were blocked in 5% milk diluted in TBST, 0.1% Tween 20 diluted in TBS (50mM Tris-HCl,pH7.5, 150mM NaCl) for 30 – 60 minutes before incubation with primary antibodies diluted in block overnight at 4°C. Primary antibodies were anti-PLP (N-terminus; 1:5000, Rogers, 2008) anti-GFP (JL8; 1:2000 – 5000; Clontech); anti-alpha-Tubulin (DM1A, 1:10000; Sigma), anti-Kinesin (SUK4 concentrate; DSHB, 1:2500), Armadillo (N2 7A1 concentrate; DSHB, 1:500), anti-Flag (Clone M2; Sigma, 1:2000). Blots were washed in TBST, 0.1% Tween20 diluted in TBS (50mM Tris-HCl, pH7.5, 150mM NaCl), the blots were incubated in secondary antibodies conjugated with horseradish peroxidase (HRP) diluted in TBST (1:2000 – 10000; Thermo Fisher Scientific). The blots were then washed and detected using SuperSignal West Dura Extended Duration Substrate (Life Technologies) and a ChemiDoc MP Imaging System (Bio-Rad).

### Ubiquitination Assay

S2 cells were transfected with plasmids containing GFP tagged PLP(F1-3) along with pMT-3xFLAG::Ubiquitin (Klebba, 2015). Ubiquitin expression was induced with 500mM CuSO_4_ after 10-15 minutes of transfection. Cells were harvested after 48 hours and proceeded to immunoprecipitation and wester blotting as described above.

### IP-Mass spectrometry

PLP F2-3 (aa 584 to 1811) protein was isolated from SF9 insect cells using Bac-to-Bac baculovirus expression system (Thermofisher). Coding sequence of PLP F2 -3 was cloned into pFastBac donor plasmid containing a Flag tag at the C-terminus of the insert site of PLP (Gift from Rong Liu, J. Sellers lab). The plasmid was then transformed into DH10Bac E.coli cells to allow recombination of PLP-F2-3-Flag with the Bacmid plasmid. The recombinant Bacmid was then transfected into SF9 insect cells followed by baculoviral infection for large scale PLP F2-3-Flag amplification. Insect cells expressing PLP F2-3-Flag cDNA were lysed in lysis buffer (50mM Tris-HCL,7.4, 500mM NaCl, 1mM EGTA, 50 μL 0.1M PMSF, 10 μL 5mg/ml leupeptin, 50 μL 1M DTT and 1 tablet of protease inhibitor (Roche). The lysate was sonicated and then centrifuged at 48,000 x g for 30 min at 4°C. The supernatant was then incubated with Flag resin for 2 hours at 4°C then harvested for washes in lysis buffer by centrifugation at 376 x g for 2 min. The protein was eluted using elution buffer (50mM Tris-HCL,7.4, 100mM NaCl, 1mM EGTA, Flag peptide (GenScript, 300 μg/ml). The isolated proteins were then prepared for mass spectrometry using an in-solution protein digestion kit (Thermo Scientific). In brief, the immunoprecipitate was desiccated by Speed Vac (Savant) and resuspended in 50 mM triethylamonium bicarbonate (TEAB), pH 8.0 with 8M urea. Samples were reduced in 20 mM DTT at 37°C for 1 hour, alkylated (40 mM iodoacetamide at room temperature 30 min), then quenched with DTT to 10mM. Samples were diluted in TAEB then and trypsin digested (∼0.1 µg trypsin at 37°C for 24 h). Samples were cleaned with C18 spin columns (Milipore) as directed by manufacturer prior to mass spectrometry analysis. Mass spec was performed by the NHLBI Proteomics core facility. Analysis was performed using Scaffold 4 (Proteome Software Inc.)

### Y2H

PLP interactions with E3 ligases were tested using yeast two hybrid assay as described (Galletta, 2015). In brief, cDNA sequence of PLP and E3 ligases fragments (cloning information in first section of methods) (shown in Figure 6E,F) were cloned to pDEST-pGADT7 (Rossignol, 2007) and pDEST-pGBKT7-Amp (Galletta, 2014) using Gateway cloning system (Life Technologies) and transformed into Y187 and Y2HGold strains, respectively. The transformants were cultured in either SD-Leu or SD-Trp media to select for those carrying the appropriate vector. The strains were mated by mixing Y187 and Y2HGold strains in yeast extract+peptone+dextrose (YPAD) medium overnight with shaking in a flat bottom 96 well plate. Diploids were selected by plating on SD-Leu -Trp dropout media (DDO). Diploids were replica-plated onto test plates: DDO to control for diploid grown and QDO plate (DDO −Ade, −Ura), DDOXA (DDO + Aureobasidin + X-α-Gal) and QDOXA (DOO −Ade −Trp −Ura + Aureobasidin A, + X-α-Gal) for interaction tests. Interactions were scored from test plates based on the presence of growth and the development of blue color, Autoactivation was identified by mating to strains carrying empty vectors.

### Male fertility

To test male fertility, individual males of the indicated genotypes were crossed to three virgin yw females and incubated at 25°C. The number of progeny produced by single male 12 to 20 days after crossing was scored. 10 or more males for each genotype was tested. We scored the degree of fertility based on the following criteria (Varadarajan, 2016): Fertile males produced more than 20 adult progeny; sterile males produced none. Any intermediated phenotypes with less than 20 adult progeny or delayed in development were categorized as sub-fertile males.

---

**Figure S1:**
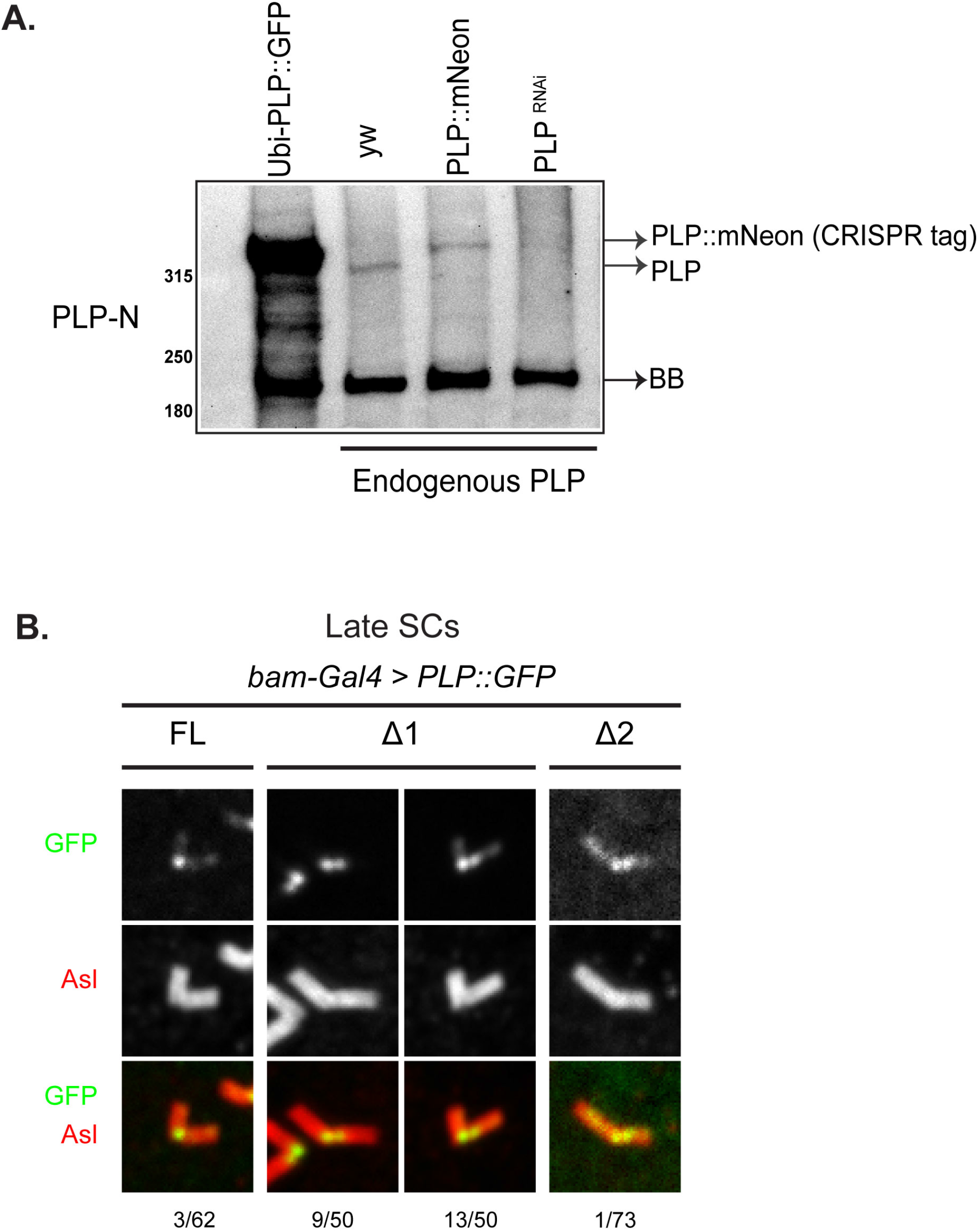
Supplement to Figure 1. **(A)** Western blot of extract from whole adult testes of the indicated genotypes. *Ubi-PLP::GFP* (lane1), *yw* (control, lane 2), *PLP::mNeon* (CRISPR knock-in, lane 3), *bam-Gal4*> *plp^RNAi^* (lane 4). Transgenic PLP::GFP expressed under a ubiquitous promoter accumulates to greater levels than endogenous PLP. The background band (BB) serves as a loading control. Sizes in kD. **(B)** Other localizations from late SCs from flies expressing the UAS-PLP deletions in Figure 1E. FL – GFP enriched at proximal end with significant GFP on more distal portions. Δ1 shows GFP at proximal end (Left), or GFP enriched at proximal end with low levels of GFP on more distal portions (Right). Δ2 shows GFP enriched at proximal end with low levels of GFP on more distal portions. Numbers indicate the frequency at which these phenotypes were observed. Bar = 1 µm

**Figure S2.**
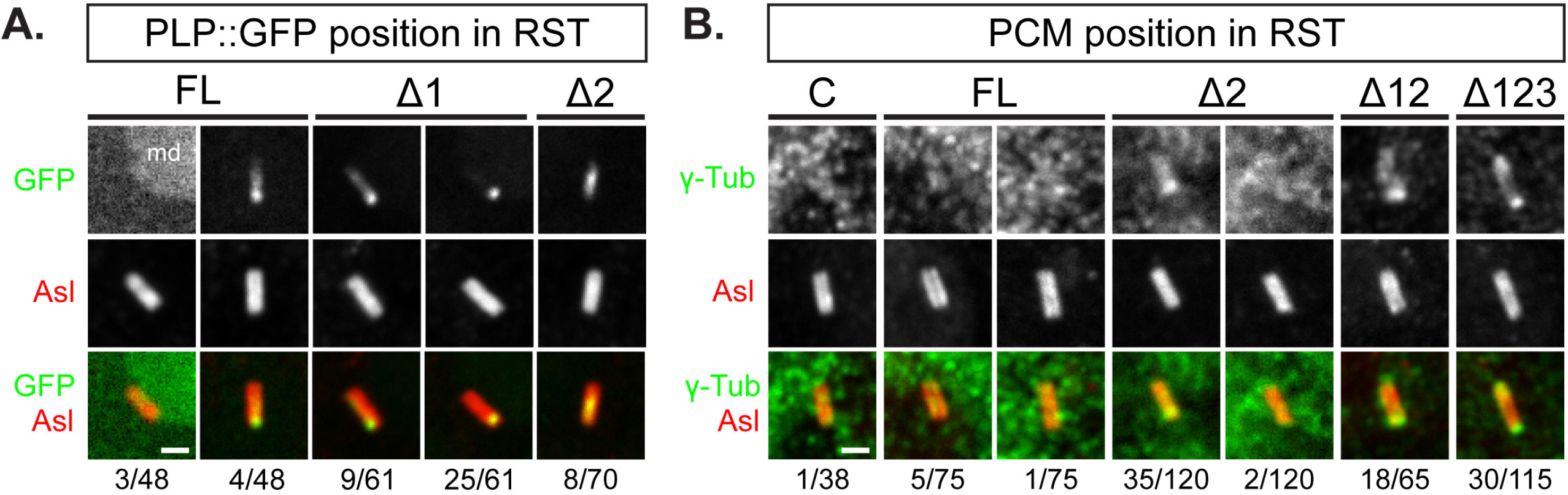
Supplement to Figure 2 showing addition localizations of PCM in spermatids with PLP deletion mutations. **(A)** Other localizations from late SCs of GFP::PLP truncations (green) and centrioles (Asl, red) in Figure 2A. Fraction of centrioles showing these GFP localization patterns are indicated below each column. FL shows no GFP observed (left), some GFP distal (right). Δ1 shows GFP along the whole length with enrichment at one end (left), GFP only at one end (right). Δ2 – GFP enriched in a patch in the middle of the centriole. Bar = 1 µm **(B)** Other localizations from late SCs of g-tubulin (PCM, green) and centrioles (Asl, red) in Figure 2B. Fraction of centrioles showing these PCM localization patterns are indicated below each column. Controle (C, *yw*) – no g-tubulin, FL – g-tubulin along the length of the centriole with no obvious enrichment (left), no g-tubulin (right). Δ2 – g-tubulin enriched on one end of the centriole (left), no g-tubulin (right). Others – g-tubulin enriched on one end of the centriole. Bar = 1 µm

**Figure S3.**
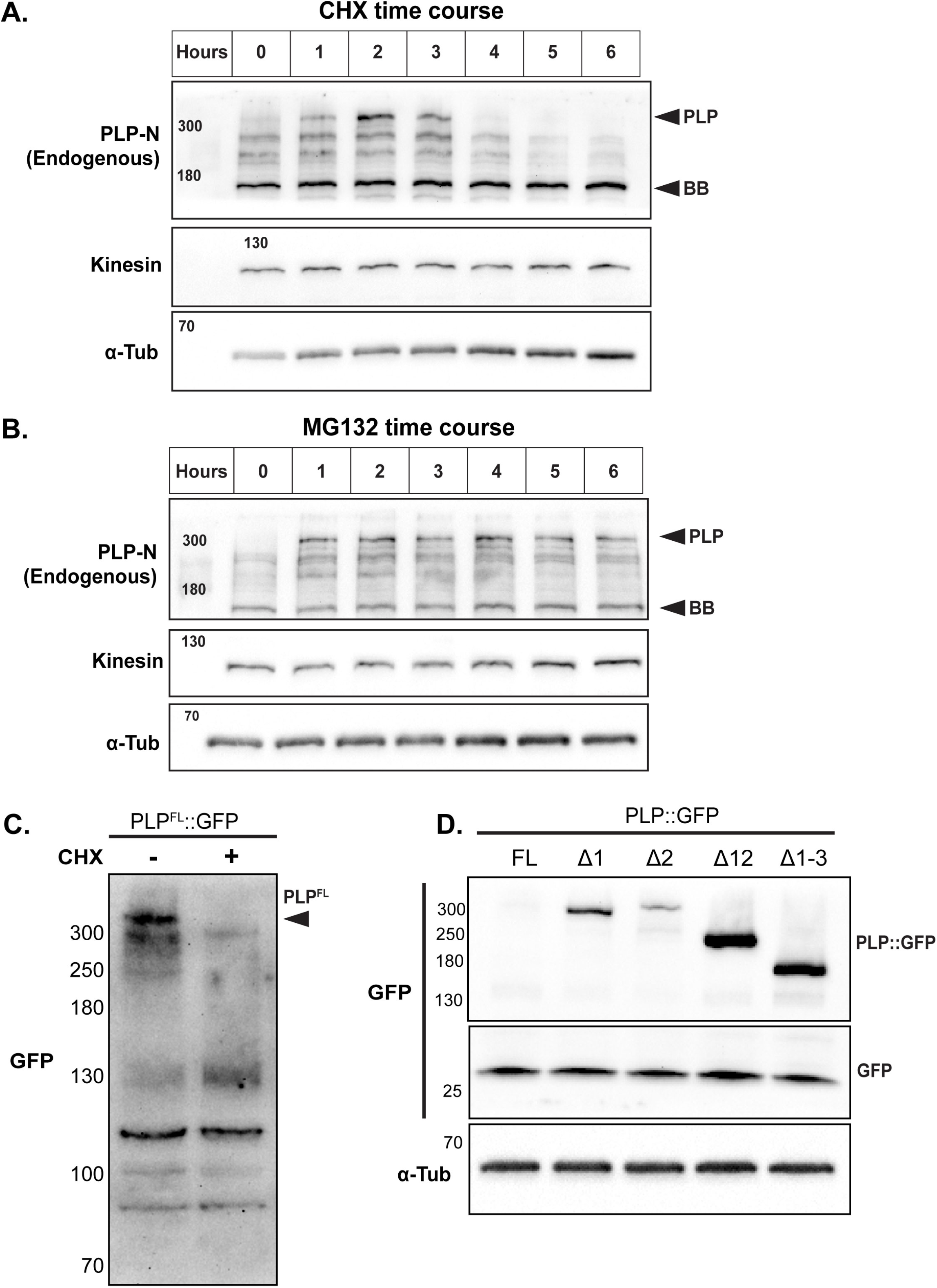
Supplement to Figure 4 showing time course of PLP levels following treatments with Cycloheximide or MG132. **(A)** Western blot of S2 cell extract showing endogenous PLP level at the indicated times following addition of cycloheximide. Kinesin, alpha-tubulin and a background band (BB) are loading controls. **(B)** Western blot of S2 cell extract showing endogenous PLP level at the indicated times following addition of MG132. Kinesin, alpha-tubulin and a background band (BB) are loading controls. **(C)** Western blot against GFP of extracts from S2 cells expressing PLP^FL^::GFP treated with DMSO or CHX for 6 hours. **(D)** Western blot showing steady-state levels of the indicated PLP::GFP constructs when expressed in S2 cells. GFP was co-transfected as a transfection control. Alpha-tubulin serves as a loading control. Sizes in kD.

**Figure S4.**
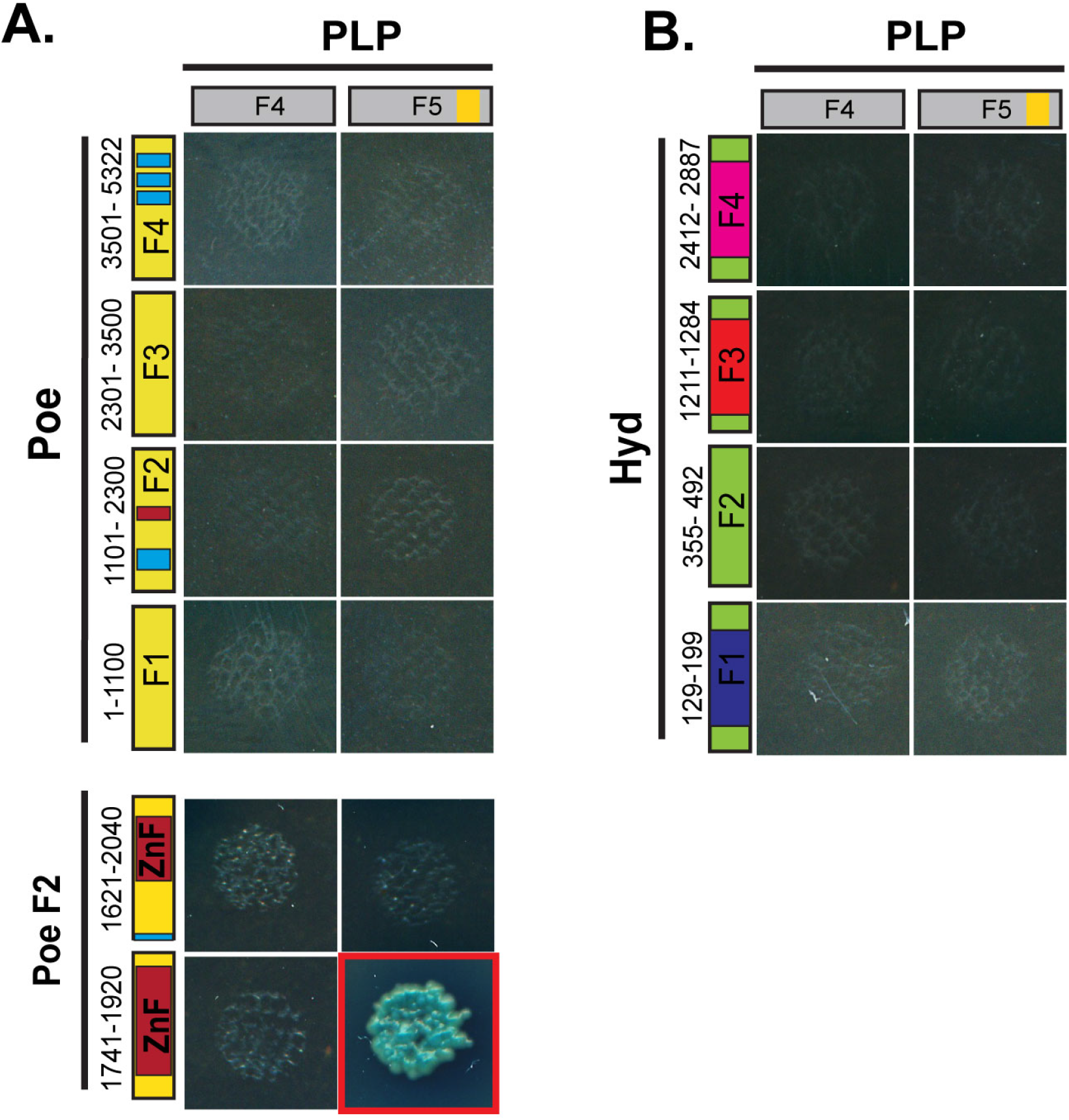
Supplement to Figure 6 showing Y2H interaction of C-terminal PLP and Poe or Hyd. Y2H interaction assay between the C-terminal fragments of PLP (F4 or F5) and all fragments of Poe (**A**) and Hyd (**B**). Presence of growth and blue color (red boxes) indicates a direct interaction between the indicated region of PLP and the E3.

## Notes

### Competing Interest Statement

The authors have declared no competing interest.

